# Spatiotemporal remodeling of cytoskeletal and junction networks during somatic cell reprogramming

**DOI:** 10.64898/2026.05.26.725436

**Authors:** Reuben Samson, Julia Kitaygorodsky, Martina Tersigni, Tulunay Tursun, Queenie Hu, William Hardy, Daniel Trcka, Hannes Röst, Jeffrey L. Wrana, Payman Samavarchi-Tehrani, Anne-Claude Gingras

**Author notes:** Correspondence to: Anne-Claude Gingras, Lunenfeld-Tanenbaum Research Institute, Sinai Health, 600 University Ave, Toronto, Ontario, Canada, M5G 1X5. These authors contributed equally to this work (Reuben Samson, Julia Kitaygorodsky).

## Abstract

Reprogramming somatic cells into induced pluripotent stem cells involves a dramatic reorganization of the cytoskeleton and junctions during the critical mesenchymal-to-epithelial transition stage. While protein abundance changes have been profiled, the spatiotemporal dynamics of protein-protein associations involving these structural components remain poorly resolved. Here, we present a time-resolved proximity proteomics resource that maps cytoskeletal and junctional remodeling across 27 baits during the early stages of reprogramming. We identified over 1100 high-confidence interactions, including many not previously reported, capturing the dynamic reorganization of cell architecture. By integrating proximity-dependent biotinylation with quantitative proteomics, we distinguished spatial relocalizations from abundance-driven effects. Dynamic redistributions of actin regulators and desmosomal proteins were observed, and a targeted short interfering RNA screen uncovered early acting structural proteins essential for colony formation. Our findings reveal adhesion and cytoskeletal maturation as structural bottlenecks in reprogramming and provide a broadly applicable framework for mapping subcellular remodeling during dynamic cell fate transitions.

## Introduction

The ectopic expression of four transcription factors (Oct4, Klf4, Myc, and Sox2; OKMS) reprograms differentiated somatic cells into induced pluripotent stem cells (iPSCs) through a highly ordered series of molecular and morphological events^1^. Time- resolved transcriptomic and proteomic studies have defined three major phases in reprogramming: initiation, maturation, and stabilization^2–8^. A key early event during initiation in mouse reprogramming is the mesenchymal-to-epithelial transition (MET), which enables the formation of compact epithelial-like colonies^2,3,9^. Proteomics profiling of a secondary mouse embryonic fibroblast (2° MEF) system that improves reprogramming efficiency revealed 2452 of 4454 proteins changed by >2-fold throughout reprogramming, with 31% (*n* = 761) of these changes occurring within the first 5 days^4^. However, the functional architecture of the cell is determined not only by which proteins are present, but also by how they are physically organized in space and time. Disruption of these interactions can derail cell state transitions, particularly if they interfere with key steps such as MET^2,3,8,10–13^. Indeed, the establishment of cell-cell adhesion during MET, classically *via* cadherin 1 (Cdh1), is indispensable for reprogramming^10,14^. Therefore, capturing the spatiotemporal changes during cell state transitions requires proteomic methods that extend beyond abundance profiling to reveal how protein environments reorganize during reprogramming.

Conventional proteomic techniques such as affinity purification–mass spectrometry (AP-MS) provide valuable insights into stable protein interactions but often fail to capture spatially defined or weak interactions^15^. This is particularly problematic in the context of morphological changes during MET, as certain interactions, such as cytoskeletal and junctional protein complexes, can be challenging to profile by AP-MS. In contrast, proximity-dependent biotinylation approaches such as BioID and its derivatives enable covalent *in vivo* labeling of proteins within ∼10–20 nm of a chosen protein of interest (bait), enabling robust recovery of proximal partners under stringent lysis conditions^16–18^; these approaches are ideally suited to studying cytoskeletal and junctional elements^19–23^. However, in general, most proximity-dependent biotinylation studies (specifically those that profile dozens of baits) have been performed in a single condition, or in conditions where proteome changes are expected to be minimal^15,24^. Furthermore, many proteins fluctuate in their abundance during early reprogramming^4,8^; therefore, it is critical to incorporate abundance measurements within the same system as proximity biotinylation studies.

Here, we report a time-resolved proximity proteomics resource that uses 27 baits to capture cytoskeletal and cell junction remodeling during early reprogramming. By integrating proximity and protein expression data, we identified more than 1100 high-confidence interactions and developed a statistical framework to distinguish spatial relocalization- and abundance-driven effects. This analysis revealed the dynamic repositioning of actin regulators and desmosomal proteins, and a functional short interfering (si)RNA screen identified structural components critical for survival and establishing early reprogramming colonies. Our findings highlight cytoskeletal remodeling and junction maturation as rate-limiting checkpoints in the MET transition and provide a broadly applicable strategy to dissect subcellular remodeling during cell fate transitions.

## Results

### Time-resolved proteomic profiling during MET in reprogramming fibroblasts

To generate a robust proteomic profile of cells during early reprogramming and to identify candidate baits for proximal profiling, we used a well-documented doxycycline (Dox)-inducible 2° MEF reprogramming line (1B)^3,25^. Derived from *piggyBac* transposition of Dox-regulated OKSM transgenes, 1B exhibits high reprogramming efficiency during the initiation phase (approximately the first 8 days) (**Supplementary Table S1**). Chimeric embryos derived from this line generated 2° MEFs that provided efficient reprogramming upon Dox induction (**Figure S1A–C**). Using a reprogramming population with ∼33% reprogramming potential, we characterized five early time points during initiation: the non-induced 2° MEF population (D0) and reprogramming days 1, 2, 4, and 6 (D1, D2, D4, and D6; **Figure 1A**). We combined isobaric peptide labeling (TMTpro16), high-pH fractionation, and liquid chromatography coupled to tandem high-resolution mass spectrometry (LC-MS/MS) to identify 7045 proteins (mapping to 6940 unique genes), with 6878 quantified across all samples (**Supplementary Table S2**). Comparisons with two previously published datasets that used similar 2° MEF reprogramming models revealed considerable overlap in identified proteins across all reported time points^4,8^ (**Figure 1B**; **Supplementary Table S2**). Furthermore, we identified an additional 1141 proteins during the early events of reprogramming that were not present in the published datasets.

**Figure 1.**
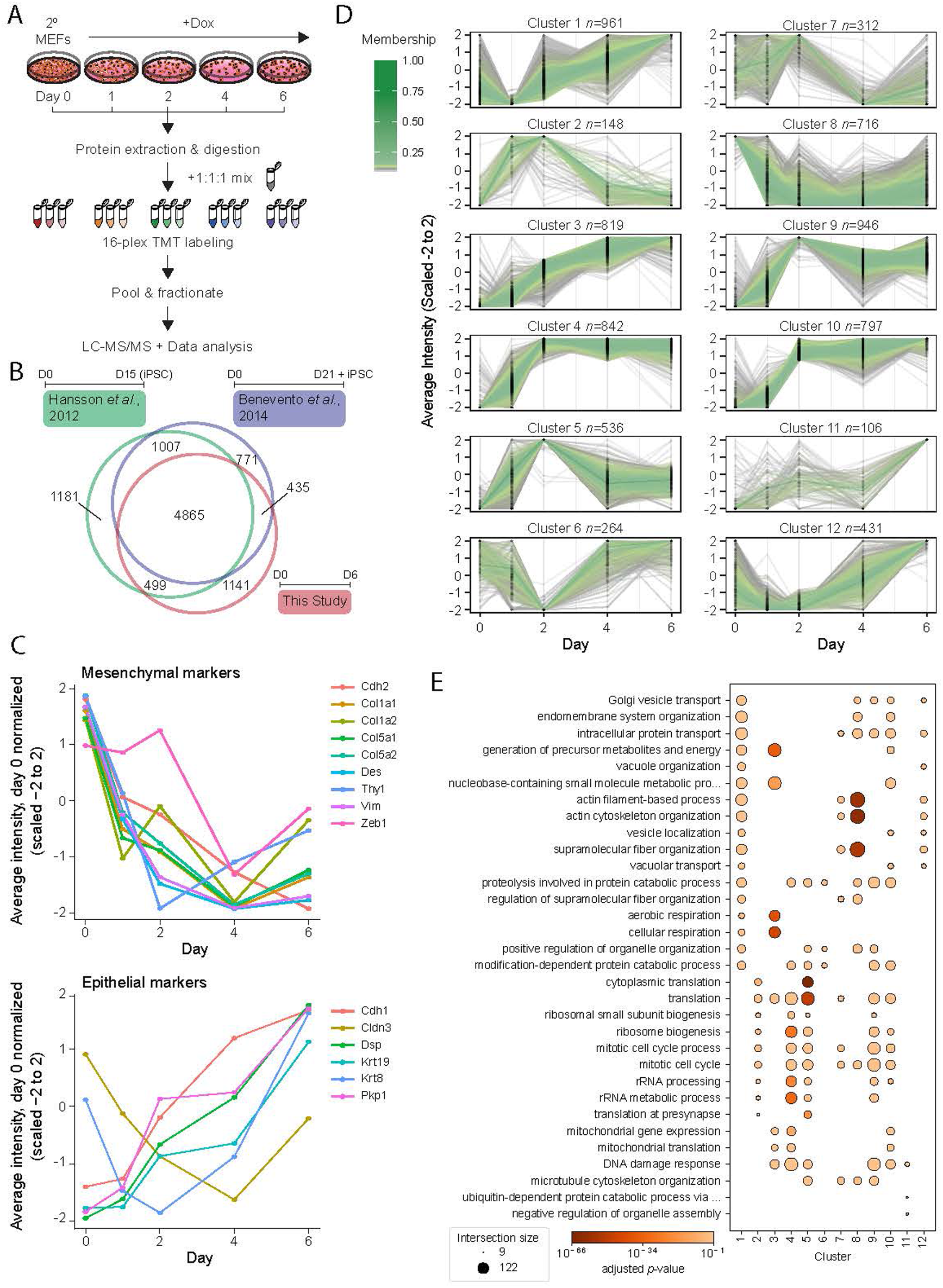
Protein expression dynamics during early reprogramming. (**A**) The total proteomics workflow: mouse embryonic fibroblasts were harvested on day 0 (non-reprogrammed) and 1, 2, 4, and 6 days after Dox induction (during reprogramming). Proteins were extracted, digested with trypsin, and labeled using tandem mass tags (TMTpro16). A pooled channel (a 1:1:1 mix of all samples) was used for normalization. Labeled peptides were combined and fractionated before LC-MS/MS analysis on an Orbitrap Fusion Lumos. Protein abundance data and peptide identification were analyzed using FragPipe software. (**B**) Venn diagram of the overlap in total identified proteins among two previous proteomics studies using a similar cellular system and this study. (**C**) Profiles of mesenchymal and epithelial markers reveal a characteristic mesenchymal-to-epithelial transition within the first 6 days. The averaged intensity value of each protein is shown, normalized to day 0 and scaled from −2 to 2. (**D**) Fuzzy c-means clustering of total proteome averaged protein intensities across days 0 to 6 of reprogramming. Protein expression profiles are scaled from −2 to 2, and membership indices are color-coded from green (high confidence, membership = 1) to gray (low confidence, membership < 0.25). (**E**) Dot plot of enriched GO biological process terms among proteins in the 12 clusters in panel D. Results are restricted to the top three GO terms per cluster (based on the lowest *p*-value; term size limited to 900).

We identified characteristic profiles of early changes in mesenchymal and epithelial markers (**Figure 1C**), indicating the successful initiation of reprogramming^3,4,26^. Our protein abundance data clustered into 12 profiles (see Methods and **Supplementary Table S2** for cluster assignment), which when analyzed for Gene Ontology (GO) term enrichment, revealed a highly coordinated change in the actin cytoskeleton (**Figure 1D** and **1E**) with clusters 1, 8, and 12 displaying an immediate drop in protein abundance within the first day, followed by a moderate return in abundance by day 6. We also identified the upregulation of proteins involved in both mitochondrial biogenesis (clusters 3, 4, and 10) and cellular metabolism and respiration (clusters 1 and 3). Notably, DNA damage response proteins were highly upregulated within our time points, consistent with previous studies^4,8^. Collectively, our data align with previously published work and confirm the robust MET that occurs in early somatic cell reprogramming.

### Proximity-dependent biotinylation of cytoskeleton and cell-cell junction resident proteins

To capture the dynamic reorganization of protein networks during MET, we applied proximity-dependent biotinylation using miniTurbo (BioID) and a Dox-inducible lentiviral delivery system^27^ (**Figure 2A**). In our 2° MEF system, Dox simultaneously induces both OKMS expression (to initiate reprogramming) and bait expression^28^. We focused on the cytoskeleton and cell-cell junctions, which undergo dramatic remodeling during the early stages of reprogramming. To systematically profile these structures, we manually curated a list of potential bait proteins localized to cytoskeletal and junctional compartments (**Supplementary Table S3**). Using our proteomic dataset, we prioritized 27 baits with minimal changes in endogenous expression over the time course (coefficient of variation < 15%; **Figure S2A** and **Supplementary Table S3**) to ensure they would serve as stable proximity sensors. Human orthologs of each bait (to enable their differentiation from mouse sequences in the mass spectrometer) were tagged with miniTurbo and delivered into 2° MEFs *via* lentivirus. Biotinylation was performed for 1 hour on days 1, 2, 4, and 6 post-Dox induction, then the cells were harvested. Peptide-level quantification of the miniTurbo ligase confirmed stable bait expression in most cases (**Figure S3A**; **Supplementary Table S3**). Across all baits and timepoints (**Supplementary Table S4**), we identified 4939 high-confidence protein-protein associations (Bayesian false discovery rate (BFDR) ≤ 1%, as assessed by SAINTexpress scoring against negative controls, see Methods), corresponding to 1161 unique preys (**Figure 2B**).

**Figure 2.**
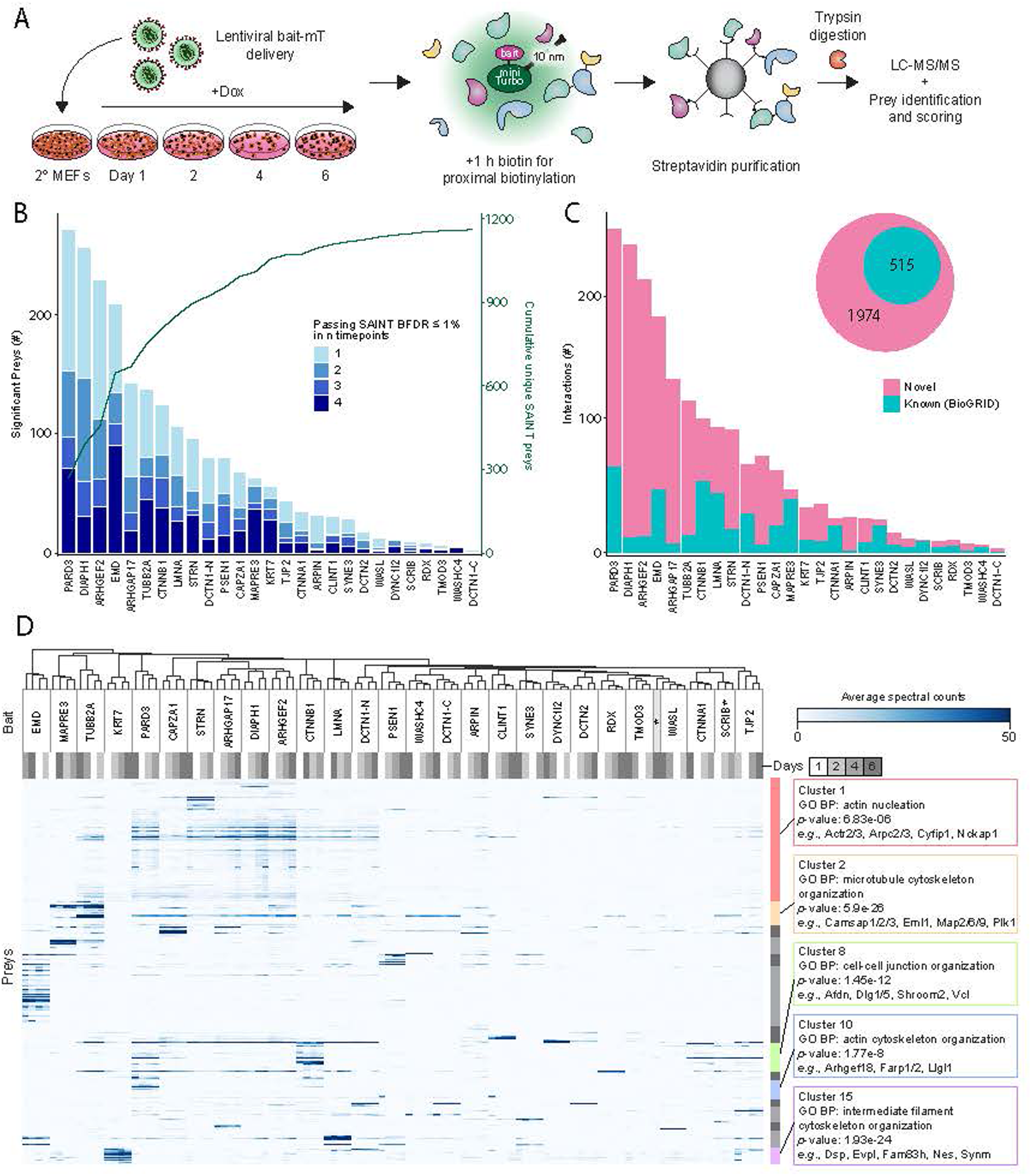
Proximity profiling of early cellular reprogramming. (**A**) The proximity-dependent biotinylation workflow. Mouse embryonic fibroblasts were transduced with lentiviral constructs encoding various cytoskeleton-related baits (human orthologs) fused to miniTurbo (mT). Upon Dox induction, both cellular reprogramming and bait expression were initiated. Biotin supplementation for 1 h at selected time points (days 1, 2, 4, and 6) enabled biotinylation of proteins proximal to the bait. Biotinylated proteins (preys) were enriched using streptavidin-conjugated beads, washed to remove non-specific proteins, and subjected to on-bead trypsin digestion. The resulting peptides were analyzed *via* LC-MS/MS using a SCIEX TripleTOF 6600, with prey identification and scoring performed using FragPipe and SAINT software prior to proximity network visualization. (**B**) Significant preys (Bayesian false discovery rates [BFDR] ≤ 1%) identified for each bait, stacked by the number of timepoints (1–4) in which preys were consistently detected. Baits are ordered along the x-axis by the total number of significant preys identified, with the cumulative unique preys plotted as a green line. (**C**) Known (*i.e.*, in BioGRID) and novel high-confidence interactors identified for each bait. (**D**) Hierarchical clustering of preys (rows) and the 27 chosen baits (columns) based on average spectral counts, following control subtraction. Clustering was performed using Pearson correlation and complete linkage, with the prey dendrogram divided into 15 clusters. Blue color saturation indicates higher average spectral counts, representing stronger proximity or abundance for a given prey-bait interaction. * = day 6 SCRIB profile.

Baits varied in the number of high-confidence preys identified, with PARD3, DIAPH1, and ARHGEF2 recovering the most across all time points (270, 254, and 227 preys, respectively), potentially reflecting their proximity to diverse protein complexes involved in cytoskeletal regulation^29,30^ and intracellular signaling^31^. Reflecting the dynamic nature of MET, most interactions were restricted to one or two timepoints, with few preys persisting across three or more. Notably, each bait identified more novel high-confidence interactions than known ones when benchmarked against human orthologs in BioGRID^32^ (**Figure 2C**; **Supplementary Table S4**), highlighting the ability of BioID to capture dynamic, weak, or spatially restricted associations.

To explore the biological processes represented in our dataset, we performed hierarchical clustering of all bait-prey interactions over time, dividing the dendrogram into 15 modules; and examined each module’s functional enrichment (**Figure 2D**, **Figure S2B** and **C**; **Supplementary Table S4**). This analysis revealed both stable and dynamic interaction profiles across baits, as well as functional insights. For instance, clusters 3 and 5 (driven by baits CAPZA1 and PSEN1) captured the dynamics of actin cytoskeleton disassembly and re-nucleation, while clusters 2 and 4 (driven by TUBB2A and MAPRE3) described microtubule regulation, with cluster 2 exhibiting a broad spatial association across BioID sensors. Clusters 8–10 highlighted dynamic interactors linked to epithelial adhesion, junction maturation, and WNT signaling, which included the baits CTNNA1, CTNNB1, PARD3, and TJP2—core components of nascent adherens and tight junctions^33,34^. These cluster-specific associations align with the known biological roles of their respective baits, reinforcing the specificity and resolution of our proximity-based approach.

### Differential associations of cytoskeletal and cell-cell junction regulators during reprogramming

BioID profiling across reprogramming revealed dynamic and stage-specific changes in the proximity landscapes of cytoskeletal and junctional regulators. For example, the CTNNA1 bait showed a day 2-specific enrichment of high-confidence actin filament binding proteins (*e.g.*, Afdn and shroom family member 2 (Shroom2)) and by day 4, a strong association with proteins involved in cell junction assembly and organization (*e.g.*, Vcl, Dlg1, Ccdc85c, and Cxadr; **Figure 3A–D**; **Supplementary Table S5**). These changes likely reflect the early reorganization of the junctional architecture and the emergence of actin-cell junction interfaces, as supported by the prominent localization of CTNNA1 to cell-cell contacts by day 6 (**Figure 3E**). Proteins involved in cellular polarity (*e.g.*, Pard3, Scrib) and Wnt pathway regulation (*e.g.*, Lzts2, Ccdc88c) also peaked in proximity at day 4 (**Figure 3A**), suggesting that epithelial polarity and signaling feedback loops are actively engaged during this phase. While CTNNA1 primarily recruited proteins associated with adherens junctions, baits such as CTNNB1 and PARD3 also captured proteins involved in tight junctions and desmosomes (**Figure 3F**; **Supplementary Table S5**).

**Figure 3.**
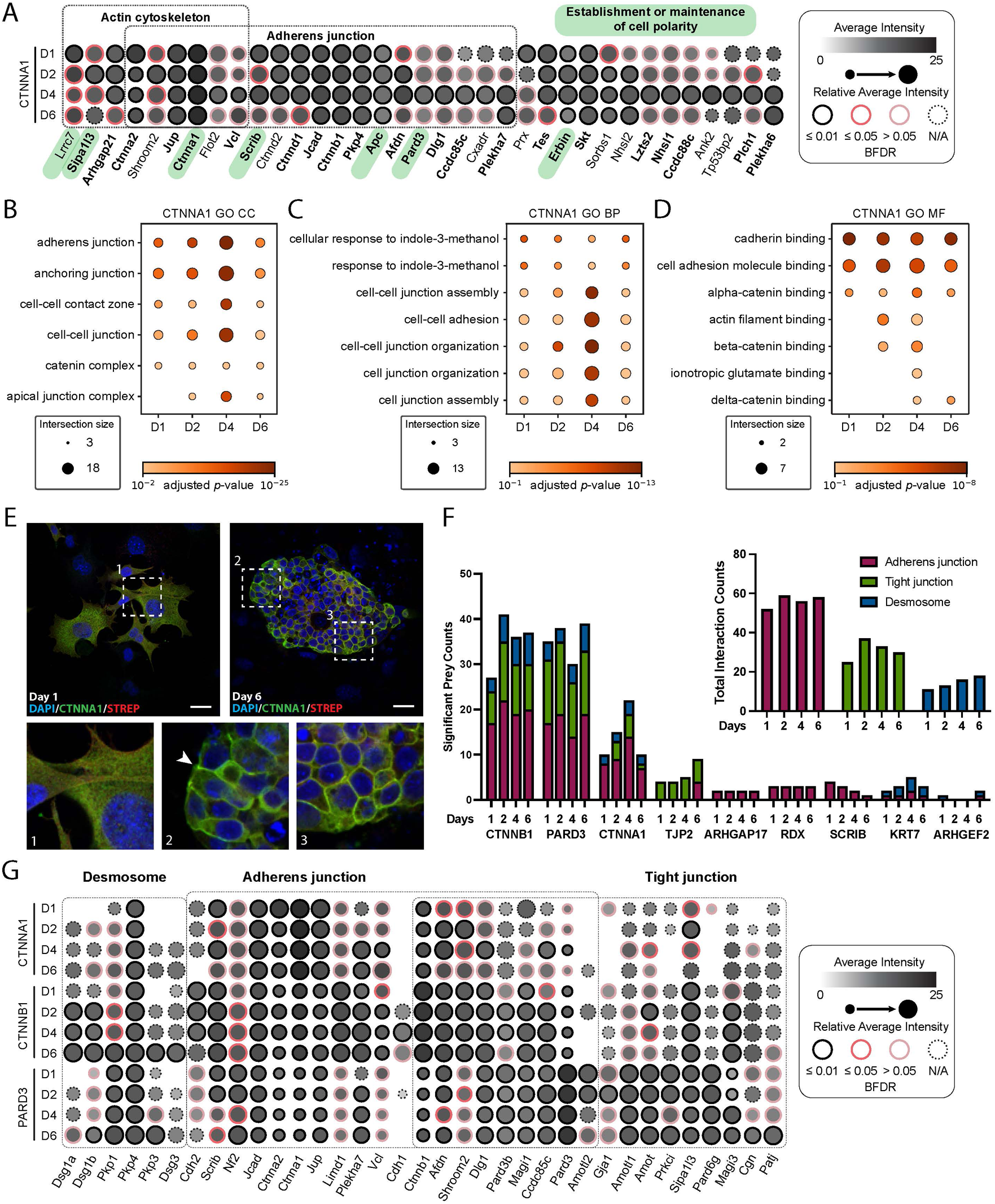
Proximity profiling reveals early cellular junction architecture remodeling. (**A**) Dot plot of preys enriched during days 1–6 for CTNNA1-miniTurbo. The average intensity is shown as a grayscale gradient (increased proximity in black) and the dot size indicates a prey’s relative intensity across days. The statistical confidence (BFDR) is shown as a color-coded ring, with N/A indicating no confidence score. (**B-D**) GO cellular compartment (CC) (**B**), biological process (BP), (**C**), and molecular function (MF) (**D**) enrichment of CTNNA1’s high-confidence (BFDR ≤ 0.01) proximity interactors during reprogramming. Only high-confidence associations for each day were used as inputs in the analyses. (**E**) CTNNA1-miniTurbo cells undergoing reprogramming were fixed and stained on days 1 and 6. Insets show magnified regions with white arrows indicating strong cell-cell contact localization. DAPI (blue), nucleus; CTNNA1 (green), miniTurbo-FLAG-tagged bait; streptavidin (red), biotinylation signal. Scale bars: 20 μm. (**F**) Stacked bar plot representing the number of significant preys localizing to the respective cellular junction components for selected baits and days. The inset shows overall counts per day. (**G**) Dot plot representing the proximity associations of selected proteins localized to the desmosome, adherens junction, and tight junction (by GO CC) with selected baits, as measured by relative average intensity.

This remodeling also aligns with recent studies describing a transient epidermal-like state during early–intermediate reprogramming^11,35^, which implicates partial keratinization as a facilitator of iPSC transition. The maturation of the desmosome and likely creation of a cornified envelope in developing colonies is supported by both our BioID data (using the keratin KRT7 as a bait; **Figure S3B**) and protein abundance profiles (**Figure S3C** and **D**; **Supplementary Table S5**), with heavy deposition of KRT7 at the colony’s periphery on day 6 (**Figure S3E**). KRT7 exhibits an increased association with structural constituents of the skin epidermis and the process of keratinization (*e.g.*, Ppl and Evpl) as early as day 2. Stabilizers and components of the desmosome (*e.g.*, Dsg1a/b, Dsg3, and Pkp3) show strong recruitment to CTNNB1 and PARD3 at later timepoints, likely indicating the late-MET formation and stabilization of the desmosome alongside other epithelial junction complexes (**Figure 3G**). More stable associations (*e.g.*, Pkp1 and Pkp4 with PARD3) and the rapid recruitment of Dsg1a and Dsg1b to CTNNB1 may indicate the nascent desmosome–adherens junction protein clusters as these components develop prior to MET^36^. Together, these findings highlight the progressive and modular assembly of epithelial junctions during colony formation and reveal the dynamic recruitment of structural regulators during MET.

### Abundance normalization to refine the identification of spatially reorganized proteins

To distinguish true spatial remodeling events from changes driven by protein abundance, we normalized our BioID proximity data against the total proteome. Of the 1161 high-confidence preys identified in our BioID dataset, 1034 (89%) were also quantified in the total proteome (**Supplementary Table S6**). Since a protein’s proximity signal can rise or fall with its total cellular abundance, we used linear modeling to detect proximity changes that were not equally proportional to abundance changes.

Based on a global linear regression analysis across all preys to evaluate the effect size of abundance on proximity labeling (**Supplementary Table 6**), a slope of 1 (the identity line) was chosen for the null hypothesis. Proteins whose measured proximity changes exceeded this expected proportionality (*i.e.,* steeper slopes) were interpreted as having stronger BioID intensity differences than could be explained by abundance alone (**Figure 4A-C**). By adopting a conservative null model (slope = 1), we minimized the likelihood of over-interpreting abundance-driven effects as spatial relocalization. This approach thus favors a stringent selection of candidate proteins for follow-up analysis.

**Figure 4.**
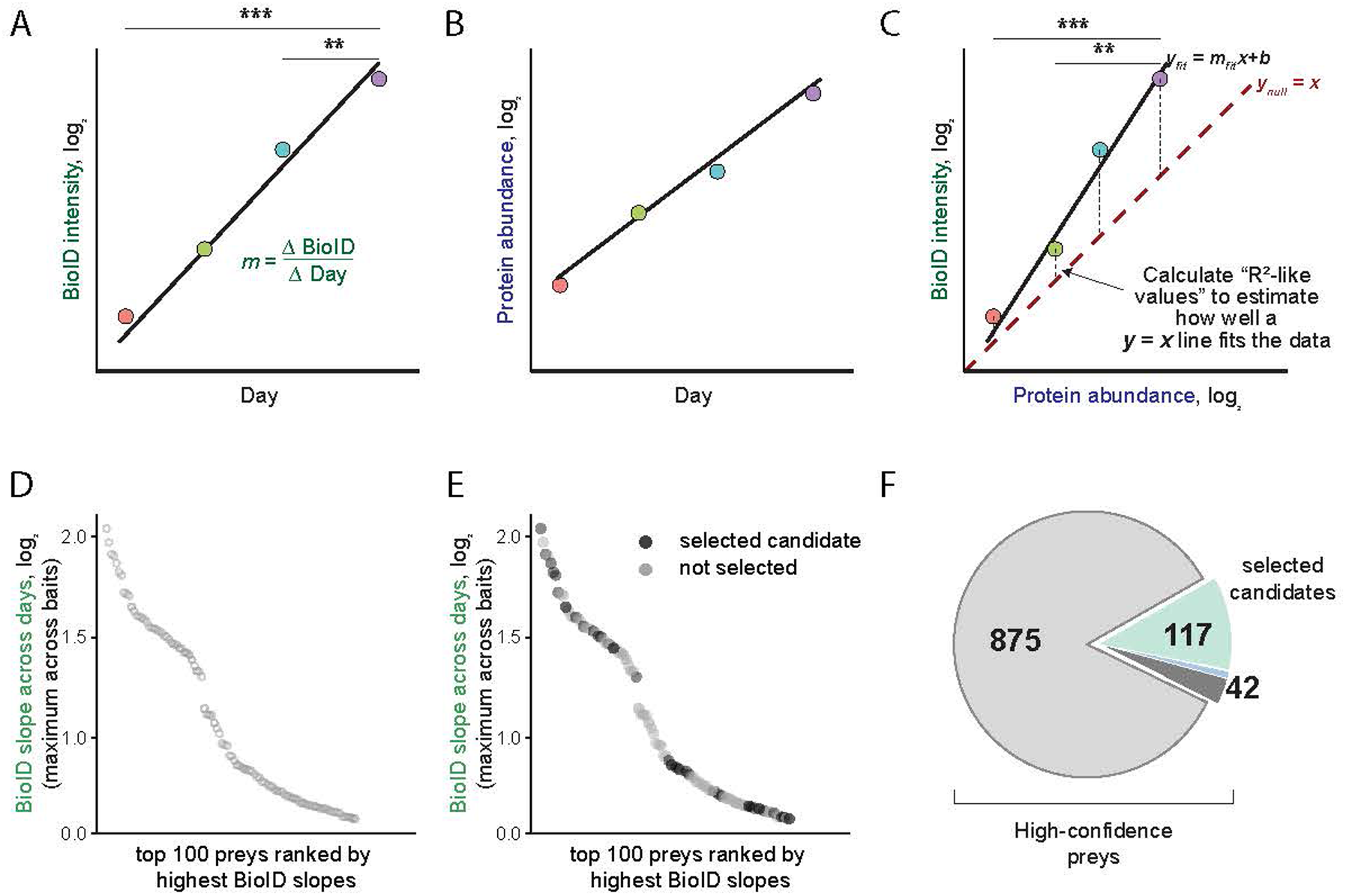
Computational method for identifying preys based on changes in abundance *versus* localization. (**A**) Example plot to determine the linear slope of the average BioID intensities for a bait-prey association over early reprogramming. (**B**) Example plot to determine the linear slope of the averaged protein abundance intensities over early reprogramming. (**C**) Schematic of the linear model used to evaluate the relationship between protein abundance data and BioID proximal association measurements for a bait-prey pair. Refer to the Methods section for details on assumptions. (**D**) The top 100 preys ranked by highest BioID slopes over time (using information only defined as in A). (**E**) Depiction of selected candidates within these top 100 preys by incorporating (B) into the linear model shown in (C). (**F**) Inclusion criteria for the final list of candidates included in the functional siRNA screen. Of the 159 selected candidates, 33 proteins were removed that were not in our siRNA collection, along with 9 of the 24 mitochondrial proteins in order to prioritize cytoskeletal components for subsequent screening.

We note that ratio compression in the total proteome quantification (resulting from the z-based TMT quantification, see Methods and Discussion) may lead to an underestimation of abundance differences across conditions. Proceeding with a conservative selection approach, therefore, helps mitigate this source of bias by avoiding false identification of abundance-related changes as proximity-specific effects.

For each bait-prey pair, we fit linear models across the four BioID time points and compared these with matched total protein abundances (see **Supplementary Table S6**). Proteins were binned by their absolute value (log_2_) slopes and R^2^-values, and each bin was scored based on model fit, magnitude of slope, reproducibility across replicates, distance to the identity line, and significance of the change in BioID proximity across days. These evaluations were combined into a mean score. From the top 2% of bins (by mean score), we selected the lowest slope and R² thresholds to define a stringent list of 205 bait-prey pairs.

To highlight the stringency of our model, only 33 of the top 100 unique preys—ranked by the magnitude of their BioID slope across time—met our criteria for spatial reorganization beyond abundance effects (**Figure 4D–E**; **Supplementary Table S7**). For example, preys of PARD3 (Anxa8 and Crym) and PSEN1 (Aars2, Irag2, Mtdh, and Srprb) exhibited significant changes in proximity across time, with minimal changes in protein abundance (**Supplementary Tables S6** and **S7**); these were consequently selected as candidates. In contrast, while two preys from KRT7’s profile (Krt17 and Krt20) also showed strong proximity shifts, these were matched by concurrent changes in abundance (and thus were not selected). As we aimed to identify preys with robust and reproducible proximity shifts independent of abundance, preys lacking temporal significance or replicate consistency (such as Atp6v0c, Cog4, Isg20l2, and Slc25a10 from DIAPH1’s profile) were also excluded.

Across all baits and preys, this conservative approach yielded a curated set of 159 unique proteins (see **Figure 4F**; **Supplementary Table S6**) whose dynamic reorganization during reprogramming is unlikely to be explained solely by expression changes.

### A functional siRNA screen identifies regulators of early colony formation

To assess the functional relevance of proteins undergoing dynamic spatial reorganization, we selected 117 of the 159 candidates (excluding mitochondrial proteins and those without available siRNAs) for a targeted loss-of-function screen for early colony formation (**Figure 4F**). Including scrambled, Oct4, and Myc siRNAs, and Dox and no-Dox controls (no siRNA), the screen contained 122 total samples (**Supplementary Table S8**). Cells were transfected with SMARTpool siRNAs in triplicate and cultured in the presence of Dox for 6 days to induce reprogramming. On day 6, colonies were fixed and stained for alkaline phosphatase (AP) to detect reprogramming and 4′,6-diamidino-2-phenylindole (DAPI) to visualize the nuclei. Colony growth and morphology were quantified using a custom automated image analysis platform that identified AP-positive colonies based on their nuclear distributions and AP signals. Replicates were averaged and ranked by AP-positive colony area (**Figure 5A**; **Supplementary Table S8**).

**Figure 5.**
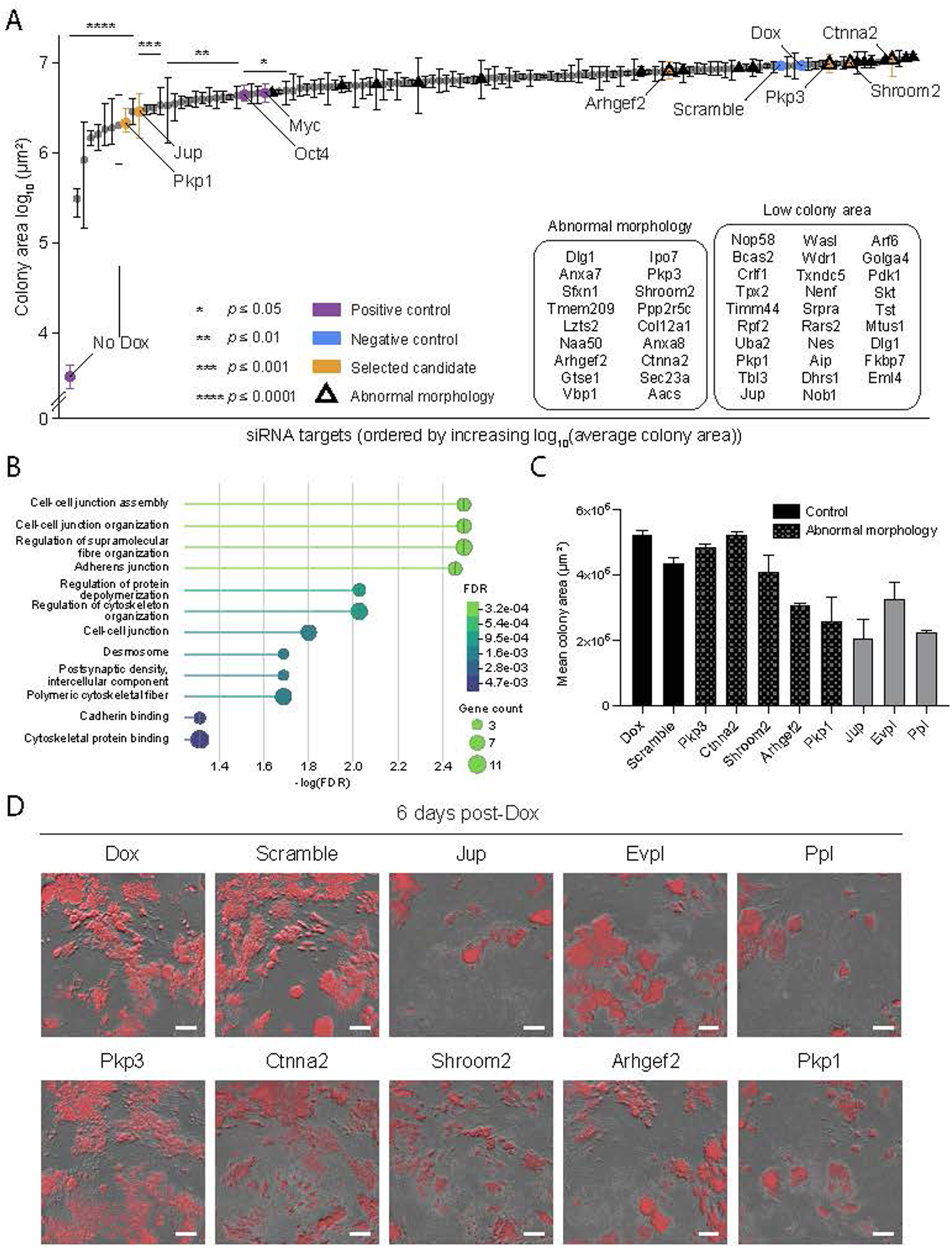
An siRNA screen reveals key regulators of reprogramming colony formation. (**A**) Rank-order plot of colony area (log_10_ of the average alkaline phosphatase (AP)-stained colony area) for siRNA-treated cells, with controls and selected candidates highlighted. siRNA constructs were introduced into cells in triplicate, and reprogramming was induced with doxycycline (Dox) for 6 days. Cells were fixed and stained for AP. The nuclei were counterstained with DAPI. Colony formation was quantified using an automated image analysis platform, which measured AP-positive, high-DAPI-density areas across four nonoverlapping fields per well. Statistical significance was assessed using ANOVA followed by Dunnett’s test for multiple comparisons. (**B**) GO enrichment analysis of targets whose siRNAs induced abnormal morphology or low colony area. (**C–D**) Individual validation of colony areas (C) and morphologies (D) for selected candidates. Scale bars: 400 μm.

In this screen, 29 siRNAs caused significantly reduced colony areas, including the Oct4 and Myc controls, which replicated known suppressive effects found in a previous study’s functional siRNA screen^3^ (**Figure 5A**). Notably, several siRNAs, including these controls (**Figure S5A**), produced colonies with abnormal morphology—despite retaining AP positivity and often displaying no significant change in AP-positive area when compared with the scrambled control—suggesting that colony area alone does not fully capture functional impairment (**Figure 5A**). Among the siRNAs that most impacted colony growth and morphology, many targeted proteins that regulate the polymerization and depolymerization of actin filaments (*e.g.*, Shroom2^37^ and catenin alpha 2 (Ctnna2)^38^ or localize to the desmosome (*e.g.*, plakophilin (Pkp)1, Pkp3, and junction plakoglobin (Jup)^39–41^; **Figure 5B; Supplementary Table S8**). This suggests that the formation of the desmosome, and likely the generation of a cornified envelope, is important for early colony formation.

To further validate this hypothesis and our screening results, we depleted six of these hits along with Evpl and Ppl, which displayed relevant BioID and abundance changes and are implicated in generating a cornified envelope, and tested their ability to impair colony formation and morphology. All candidates replicated the findings from the larger screen, and Pkp1 depletion also induced abnormal colony morphology (**Figures 5C** and **D, Figure S6A; Supplementary Table S8**). Depleting Shroom2, Ctnna2, or Arhgef2—regulators of actin cytoskeleton organization—resulted in distinctly smaller colonies that were AP-positive but had mesenchymal-like morphologies (*i.e.*, flat and 2D). Pkp1, Jup, Evpl, and Ppl knockdowns all reduced the overall colony area, with Pkp3 and Pkp1 depletion causing similar abnormal colony morphologies with loose colony boundaries. Together, these findings suggest that desmosome maturation and stabilization, and actin cytoskeleton reorganization are important steps in early colony formation.

### Early regulators of colony structure are essential for successful reprogramming

Although all eight siRNAs impaired colony formation in terms of overall growth area, morphology, or both, four (targeting Pkp3, Ctnna2, Shroom2, and Arhgef2) showed no significant differences in mean colony area at day 6 compared with negative controls. As the initiation-maturation phase transition occurs between days 6 and 8 in this model^3^, we investigated whether knockdown of these candidates would accelerate MET or impact overall colony survival post-MET. Despite some knockdowns displaying high colony areas at day 6, none displayed improved survival after Dox removal at day 9 (**Supplementary Table S9**). Instead, most colonies failed to persist beyond this point, in contrast to controls (**Figure S6C**). To evaluate their potential to form stable, Dox-independent colonies, we extended Dox treatment to day 12, enabling expression of early pluripotency markers such as *Nanog* and *Sall4*, which are required for stable acquisition of pluripotency^42,43^ (**Figure 6A; Supplementary Table S9**).

**Figure 6.**
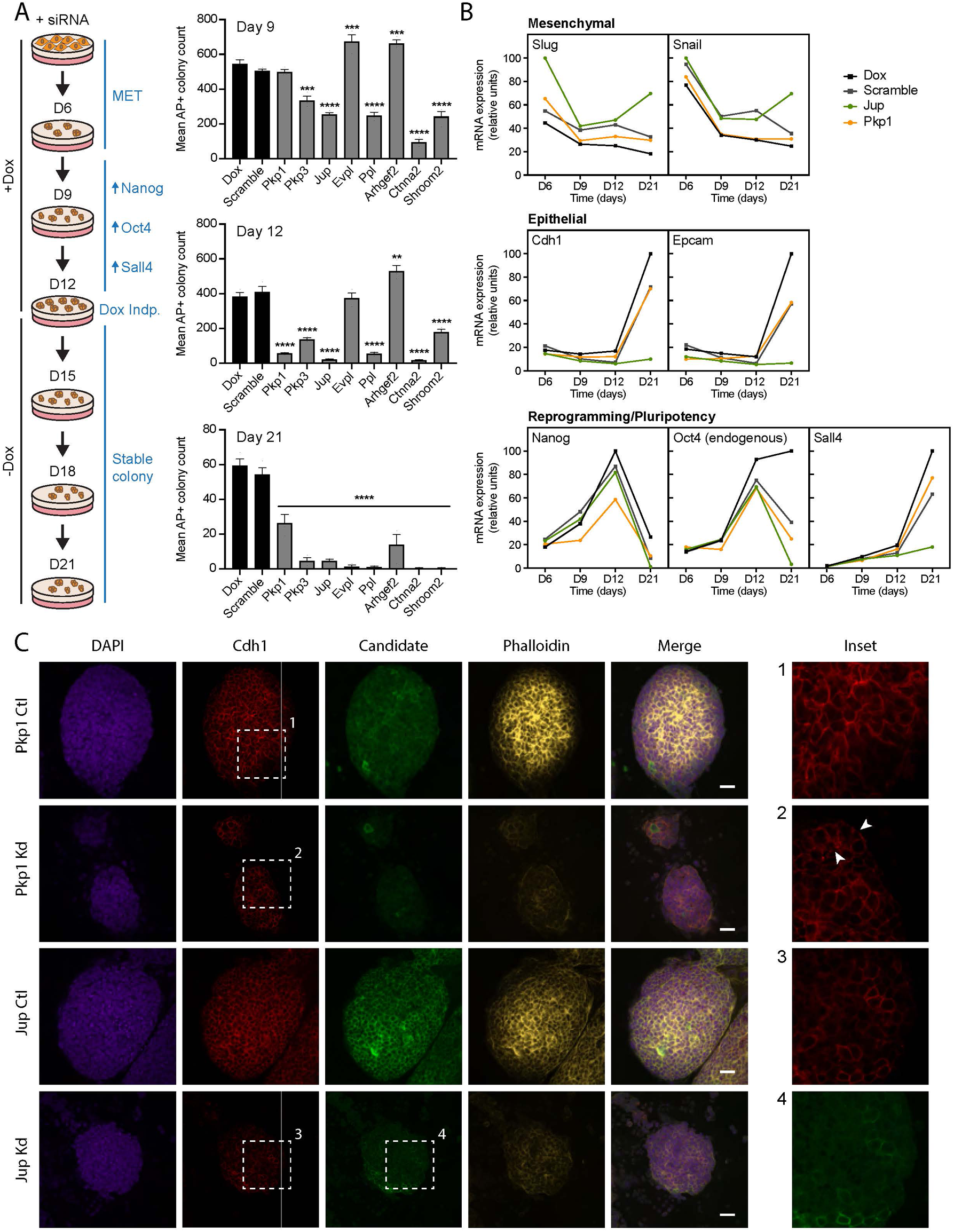
Candidate knockdown prevents transition into fully reprogrammed colonies. (**A**) Alkaline phosphatase (AP)-positive colonies were counted on day 9 and day 12 (with doxycycline (Dox) supplementation) and day 21 (post-Dox removal at day 12) following siRNA knockdown of selected candidates. Cells were split (1:60) onto feeder mouse embryonic fibroblasts every 3 days starting on day 6. The blue schematic outlines the typical steps in reprogramming; Dox indp., beginning of doxycycline independence. Asterisks indicate one-way ANOVA adjusted *p*-values (** = <0.01, *** = <0.001, **** = <0.0001). (**B**) RT-qPCR was performed to measure mesenchymal, epithelial, and reprogramming/pluripotency markers on days 6, 9, 12, and 21 upon siRNA knockdown of Jup and Pkp1. (see **Figure S7** for data from all candidates shown in A). Transcript levels are normalized to beta-actin. (**C**) Immunofluorescence of reprogramming colonies on day 6 after transfection of a scrambled siRNA (Ctl samples) or siRNAs targeting Pkp1 and Jup (Kd samples). Insets provide magnified views of cell-cell junction formation (Cdh1) and/or candidate localization. White arrows indicate fragmented Cdh1 staining. DAPI (blue), nucleus; Cdh1 (red), cadherin 1; candidate (green), Pkp1 or Jup; phalloidin (yellow), F-actin. Scale bars: 20 μm.

Colony counts varied at day 9, with Jup, Ppl, Ctnna2, and Shroom2 depletion resulting in significant losses relative to controls. By day 12, all knockdowns except Evpl and Arhgef2 displayed reduced colony numbers. Following Dox withdrawal after day 12, all knockdowns induced a marked collapse in colony numbers, suggesting that these perturbations compromised the ability to stabilize the reprogramming trajectory. Together, these results indicate that most proteins tested play important roles during the early stages of reprogramming—particularly within the first 6–9 days—contributing to both colony formation and the maintenance of epithelial colony integrity through MET.

Exploring MET and reprogramming/pluripotency marker expression changes by reverse transcription-quantitative polymerase chain reaction (RT-qPCR) provided further insight into how depleting these candidates affects early colony formation (**Figures 6B** and **S7; Supplementary Table S9**). Like the negative controls, all knockdowns exhibited characteristic decreases in mesenchymal marker expression (*e.g.*, *Slug*, *Snail*) within the first 9 days of Dox treatment. However, by day 21, all except for Pkp1 (which largely resembled the controls), had returned to increased levels of at least one of *Slug* and *Snail*, suggesting that MET was not maintained. While epithelial marker expression increased in the scrambled and Dox controls by day 21, this was not the case with six of the knockdowns, confirming the failure to complete MET. Pkp1 again resembled the controls, and Arhgef2 displayed an intermediate stage.

Among the knockdowns, Pkp1, Pkp3, and Jup showed increased levels (at least 50% of control levels) of the reprogramming/pluripotency markers *Nanog* and (endogenous) *Oct4* by day 12, indicating some level of pluripotency establishment (**Figures 6B** and **S7**). Although there was a modest loss in *Snail* and *Slug* expression, Arhgef2 depletion did not yield strong expression of pluripotency markers by day 12, but did express both epithelial and reprogramming/pluripotency markers (*Oct4* and *Sall4*) by day 21. This delay may be due to dysregulated and poorly organized actin cytoskeletons in most budding colonies in late-MET onwards (days 6–12), with those managing to survive to day 21 driving the marker profile of Arhgef2 depletion (as the high colony counts seen on days 9 to 12 were minimal by day 21).

Considering that Pkp1 and Jup knockdowns both showed significant mean colony area losses within the first 6 days of reprogramming, but were able to exhibit successful MET based on marker expression levels (in the case of Pkp1) or establishment of pluripotency/reprogramming marker expression (for both Pkp1 and Jup), we examined how their loss affected colony formation through immunofluorescence. Pkp1 knockdown colonies exhibited weak and disrupted cortical actin staining and organization, with fragmented Cdh1 staining at cell-cell junctions on day 6 (**Figure 6C**). Jup knockdown colonies exhibited reduced Cdh1 (coinciding with the RT-qPCR results) and cortical actin staining throughout, with strong Cdh1 deposition only where Jup was localized. This emphasizes the likely role of Jup in recruiting and stabilizing adherens junction proteins^44^, such as Cdh1, during MET. Moreover, the absent or poor stability of epithelial junction organization (as indicated by low or disrupted Cdh1 expression) within knockdowns of both Pkp1 and Jup may indicate an impaired feedback mechanism (*e.g.*, *via* a Cdh1-mediated signaling event) that supports epithelialization and the pluripotency network^12^. Overall, our data highlight a series of proteins that underpin the structural transitions required for MET, which are indispensable for the stable formation of reprogrammed colonies.

## Discussion

Inducing pluripotency in differentiated somatic cells *via* MET involves a highly coordinated remodeling of the cell’s molecular landscape^2,3^. Although time-resolved quantitative proteomics have extensively characterized protein abundance changes across this critical transition^4,8^, the spatiotemporal organization of the protein-protein associations driving this remodeling remains less defined. In this study, we combined quantitative proteomics with time-resolved BioID proximity mapping using 27 cytoskeletal and cell junction baits across four early reprogramming time points. Integrating protein abundance and spatial proximity data with statistical modelling allowed us to identify 117 high-confidence proteins exhibiting significant spatial reorganization independent of changes in their abundance.

An siRNA screen of these hits revealed critical roles for cytoskeletal and junctional proteins in the early stages of colony formation, as depleting proteins such as Jup, Pkp3, Ctnna2, and Shroom2 significantly impacted colony morphology and structure within the first 6 to 9 days, suggesting indispensable roles during early reprogramming. These proteins are central to cytoskeletal remodeling and junctional stability, and their depletion disrupted actin-junction integrity^37,38^, desmosome maturation, and precursor complex formation^40,45,46^.

Interestingly, depleting Evpl and Ppl—primarily known for their roles in cornified envelope formation downstream of desmosomes^47^—induced minimal phenotypes in early reprogramming but led to substantial colony loss in later stages. This temporal specificity likely reflects roles in desmosome maturation and/or epithelial stabilization after initial MET. Conversely, Arhgef2 knockdown permitted early colony formation (including AP positivity) but ultimately compromised late-stage colony integrity. Arhgef2 regulates Rho/Rac-dependent cortical actin formation^48^ and cell polarity^29^. This suggests that its early absence promotes actin depolymerization—a key feature of early MET—but impairs the formation of the stable cortical actin network necessary for junctional maturation and colony maintenance.

Our findings underscore the crucial roles of Pkp1 and Jup in the assembly and stabilization of epithelial junctions. Pkp1 is essential for desmosome formation^39,49,50^ and Jup plays roles in both desmosomes and adherens junctions^41,44,51^. Despite apparent progression through MET and pluripotency marker activation, Pkp1-depleted colonies exhibited disrupted cortical actin and Cdh1 staining patterns. This suggests a key role for Pkp1 in stabilizing nascent desmosomes, and possibly orchestrating the spatial segregation and maturation of adherens junction and desmosomal proteins^36^ in parallel with cytoskeletal remodeling^52^. Depleting Pkp1 or Jup ultimately led to most colonies failing to achieve stable pluripotency, highlighting the critical roles of transient desmosome formation^11^ and junctional complex maturation in the successful growth of reprogramming colonies.

We recognize that there are limitations to our approaches. One is the ratio compression inherent to MS-based TMT quantification^53^. Co-isolation of precursors can generate background signals across reporter channels, underestimating fold changes and creating bias toward conservative interpretations. Alternative reporter-free MS-based measurements, such as data-independent acquisition^54^ (DIA), should provide a good parallel to isobaric-labeling methodologies^55^. DIA-based approaches are label-free (and therefore are not hindered by ratio compression) and offer minimal sample processing while maintaining reproducible and comprehensive proteome quantification.

Additionally, our use of R^2^ values to model proximity-abundance trends may overlook biologically meaningful outliers. For instance, Phb2 and Cog5 were excluded because of poor R^2^ fits, despite displaying interesting proximity dynamics. Since R^2^ measures variance along the y-axis, it penalizes vertical data patterns; such cases may be better assessed using orthogonal R^2^ or regression to a vertical axis. Notably, these cases often involved missing values (which were replaced with zero values), which further reduced confidence in those estimates. Future modeling could benefit from alternative strategies such as robust regression, orthogonal distance regression, or imputation-based confidence scoring.

Temporal resolution was another constraint. The need to passage reprogramming cells every 2–3 days post-initiation phase^28^ limited the expansion of reprogramming BioID into later stages. As larger colonies are broken up during resuspension, this passaging could affect the natural re-arrangement of the cellular architecture as cells recover, making certain time points unavailable for BioID profiling. Therefore, further studies should carefully examine the time points used for profiling in relation to the need to perturb the cells from their natural growth stage.

In conclusion, our integrative proteomic and spatial mapping approach identified critical structural proteins governing successful MET and colony stabilization during early somatic cell reprogramming. These findings emphasize that structural and junctional integrity are as critical as biochemical signaling for pluripotency induction, revealing novel mechanistic insights into cell fate transitions and opportunities to further optimize reprogramming methodologies.

## Methods

### Chimera aggregation, secondary reprogramming MEF isolation, and cell culture

Chimera aggregation and MEF isolation were performed as described previously^28^. Briefly, chimeras were produced by aggregating 1B cell line iPS cell clumps with Hsd:ICR(CD-1) embryos and implanting them into pseudo-pregnant females. Harvested embryos (14.5 days post-conception ROSA26-reverse tetracycline-controlled transactivator (rtTA)-IRES-GFP from t(ROSA)26Sortm1.1(rtTA,EGFP)Nagy) were decapitated, eviscerated, dissociated with 0.25% trypsin and 50 mM ethylenediaminetetraacetic acid (EDTA), and plated in MEF media (Dulbecco’s modified Eagle’s medium (DMEM) high glucose, 10% fetal bovine serum (FBS), 1% MEM non-essential amino acids, 0.5% Pen/Strep) on separate 0.1% gelatin-coated plates. Each plated embryo was expanded 4-fold prior to pooling and freezing. To create a pool of cells with significant reprogramming potential for proteomics experiments, the chimeric contributions for each embryo were assessed by flow cytometry on a Beckman Coulter Gallios flow cytometer. The analysis workflow included scatter pulse discrimination to remove aggregates, viability gating to remove dead cells, and gating to distinguish true green fluorescent protein (GFP)-positive (*i.e*., OKMS-positive) cells from the autofluorescent background (CD-1 MEFs were used as negative controls). 2° MEF populations were then pooled to generate cell batches with varying reprogramming potential (see **Figure S1** and **Supplementary Table S1**) and flash frozen in liquid N_2_ for further experiments.

For reprogramming, cells were cultured in mES media (DMEM high glucose, 15% FBS, 0.5% Pen/Strep, 0.5% GlutaMAX, 1% MEM non-essential amino acids, 1:150,000 beta-mercaptoethanol, and leukemia inhibiting factor at 1000 U per 500 mL (obtained from the Stem cell Core Facility at the Lunenfeld-Tanenbaum Research Institute (LTRI)) supplemented with 1 µg/mL Dox (to induce OKMS and bait expression) on 0.1% gelatin-coated plates. The medium was changed daily.

### Total proteome sample preparation

2° MEFs (day 0) and reprogramming populations (days 1, 2, 4, and 6) were harvested in triplicate and centrifuged at 500 × *g* for 5 min at 4°C prior to snap-freezing. Cell lysis was performed using a 1:10 (pellet weight/lysis buffer) volume of modified radioimmunoprecipitation (modRIPA) buffer (50 mM Tris-HCl, pH 7.4, 150 mM NaCl, 1 mM ethylenebis(oxyethylenenitrilo)tetraacetic acid, 0.5 mM EDTA, 1 mM MgCl_2_, 1% NP-40, 0.1% sodium dodecyl sulfate (SDS), 0.4% sodium deoxycholate, 1 mM phenylmethylsulfonyl fluoride, and 1× Protease Inhibitor Cocktail [Sigma-Aldrich, Cat# P8340]). Lysates were sonicated for 15 sec (5 sec on, 3 sec off for three cycles) at 30% amplitude on a Q500 Sonicator with a 1/8-inch Microtip (QSonica, Newtown, Connecticut, USA, Cat# 4422). Subsequently, 1:1000 volumes of TurboNuclease (BioVision Inc., Milpitas, CA, USA, Cat# 9207) and RNase A (Bio Basic, Markham, ON, Canada, Cat# RB0473) were added, and samples were rotated at 4°C for 15 min. The SDS concentration was increased to 5% (by adding 10% SDS) and samples were rotated at room temperature (RT) for a further 15 min. Samples were centrifuged at 15,000 × *g* for 15 min at RT and the supernatants were transferred to new tubes. The protein concentration of each sample was quantified using the Bio-RAD DC Protein Assay Kit per the manufacturer’s guidelines. A 100 μg aliquot of each sample was added to a new tube and modRIPA was added to a 32 μL total volume prior to reduction with 8 μL 100 mM dithiothreitol (20 mM final) at 56°C for 30 min followed by alkylation with 10 μL 200 mM iodoacetamide (40 mM final) at RT in the dark for 30 min. Samples were clarified for 8 min at 13,000 × *g* and transferred to new tubes, then acidified with 5.5 μL 12% phosphoric acid prior to S-Trap (ProtiFi) digestion. To each trapped sample, we added 20 µL trypsin in digestion buffer (250 ng/μL in 50 mM tetraethylammonium bromide (TEAB)), and the mixture was incubated for 2 h at 45°C. Peptides were eluted sequentially with 40 μL each of 50 mM TEAB and 0.2% formic acid at 4,000 × *g* for 30 s. A final elution with 35 μL 50% acetonitrile containing 0.2% formic acid was performed prior to lyophilization. Samples were resuspended in 100 μL 0.2% formic acid, and 5 µL was used for peptide quantification using the Pierce Quantitative Colorimetric Peptide Assay kit (Thermo Fisher Scientific) following the manufacturer’s recommendations. Aliquots of each sample (20 μg) were re-lyophilized, resuspended in 20 μL 100 mM HEPES pH 8.5, and labeled with 8 μL Tandem Mass Tag reagents (160 μg TMTpro16, in acetonitrile) for 1 h at RT. After quenching with 8 μL 5% hydroxylamine for 15 min, 18 μL (10 μg) of each sample was lyophilized, resuspended in 30 μL 0.1% trifluoroacetic acid, and pooled prior to fractionation using a Pierce High pH Reversed-Phase Fractionation kit (Thermo Fisher Scientific) into eight fractions with an increasing acetonitrile gradient (10–50%). Fractions were lyophilized prior to resuspension in 5% formic acid.

### Total proteome mass spectrometry

Fractions were analyzed in data-dependent acquisition (DDA) mode on an Orbitrap Fusion Lumos Tribrid MS (Thermo Fisher Scientific) connected to a 425 Nano-HPLC system. The MS1 scan was over a mass range of 400–1500 *m/z*, with a 50 ms accumulation time, Orbitrap resolution of 240,000, 60% RF lens, and 2600 V. MS/MS scans had an 86 ms accumulation time, 35% HCD collision energy for ions charged 2+ to 7+, and an automatic gain control target of 5e4 (isolated using an Orbitrap resolution of 50,000 Da [dynamic exclusion 12 s]). The total cycle time was 3 s.

### Total proteome data analysis

Total proteome data were converted to mzML format using MSConvert (version 3.0.22290) with “Peak Picking” filter set to “Vendor” and 64-bit encoding. These mzML files were searched on the FragPipe computational platform (offline from our local interaction proteomics laboratory information management system ProHits^56^). Files were searched against a mouse protein database from UniProt (downloaded January 22, 2024) containing only reviewed sequences and supplemented with contaminants and reversed sequences acting as decoys. Default parameters were used with the TMT16 workflow.

The entire total proteome dataset was deposited in ProteomeXchange *via* partner MassIVE (massive.ucsd.edu) and assigned accession numbers PXD075628 and MSV000101129.

### Total proteome clustering

Unsupervised clustering of total proteome data was performed by fuzzy c-means using the e1071 package^57^ in R (version 1.7-6), with a Manhattan distance method and 3.5 fuzziness parameter. A range of 3 to 15 centers was assessed for data clustering, and we selected 12 clusters based on the mean silhouette score across ten random seeds. Random seed 713532 had the highest silhouette score with 12 clusters and was maintained for consistent results. Proteins were attributed to a cluster based on their highest membership value and plotted as line graphs (color-coded by membership value) using the ggplot2^58^ R package (version 3.3.5).

### Generation of lentiviral BioID expression constructs

Gateway-compatible entry clones (sourced from the Mammalian Gene Collection or Network Biology Collaborative Centre Human ORFeome V8.1 clone collections or generated *de novo* from cDNA) for all genes were transferred into the pSTV2-miniTurbo-FLAG Gateway-compatible lentivirus destination vector (N- or C-terminal tagged; see **Supplementary Table S3**) using standard LR-Gateway cloning. A control vector expressing enhanced (E)GFP (eGFP-mT) was generated to allow for background subtraction. The N-terminal pSTV2-miniTurbo-FLAG destination vector was used to create an enzyme only (N-mT) control. Non-transduced cells were also used as an empty control. All vectors were verified by restriction enzyme digestion (BsrGI) and partial forward and reverse sequencing to confirm insertion of the open reading frame.

### Lentiviral production, cell pool generation, and BioID

HEK293TN cells (System Biosciences) were used for virus production, as described in detail elsewhere^27,28^. Briefly, 1.3 µg psPAX2, 0.8 µg VSV-G, and 1.3 µg transfer vectors (pSTV2) were transfected into HEK293TN cells (at 80% confluence in a 6-well plate) using jetPRIME reagent per the manufacturer’s recommendations. After 6 h, the medium was replaced with 3 mL virus production media (DMEM high glucose, 5% heat-inactivated FBS, 1% Pen/Strep). Virus was harvested 48 h post-transfection, cleared by centrifugation (500 × *g*), filtered through a 0.45-µm filter, and frozen in aliquots at −80°C. The viral titer was estimated by co-infecting HeLa cells (at 40% confluence in a 24-well plate) with 25–100 µL of the pSTV2-miniTurbo-FLAG-tagged baits (or controls) and an rtTA viral supernatant. The next day, the medium was replaced with medium containing 1 µg/mL Dox for 24 h, supplemented with biotin (40 µM final) 1 h before harvest. Cells were subsequently fixed and analyzed by immunofluorescence (staining for the FLAG epitope, biotinylated proteins, and nuclei; see the *Immunofluorescence* section below) to determine the optimal amount of supernatant to yield a 75–85% infection rate. These results provide the infection parameters for MEFs, as we observed no significant difference in the transduction potential of HeLa cells and the MEFs used in this study.

For all BioID experiments, 2° MEFs (∼33% reprogramming potential) at approximately 60% density in single 10-cm gelatin-coated dishes were infected in duplicates with the amount of virus determined to yield 75–85% infection. For most lentiviral baits, this corresponded to 500–900 µL viral supernatant. To account for proliferation over time, cells were differentially split (described previously^28^) into gelatin-coated 10-cm plates to yield sufficient material for multiple time points of reprogramming BioID and downstream MS analysis. An aliquot of each cell suspension was also plated onto a gelatin-coated coverslip for immunofluorescence. Reprogramming was induced using 1 µg/mL Dox in reprogramming media, which was supplemented with biotin (40 µM final) 1 h before gentle harvesting with cold trypsin to generate ∼60 mg cell pellets (equivalent to one confluent 10-cm plate) for each time point and biological replicate, which were flash frozen.

### Streptavidin capture and digestion

Streptavidin affinity purification was carried out as described previously^28^. Briefly, and as with the total proteome lysates, cells were lysed in a 1:10 (pellet weight/lysis buffer) volume of modRIPA buffer prior to sonication and incubation with TurboNuclease and RNase A. Next, the SDS concentration was increased to 0.5% (by adding 10% SDS) and the samples were rotated at 4°C for a further 5 min. Samples were centrifuged at 15,000 × *g* for 15 min and the supernatant was used for biotinylated protein capture using a 15 µL bead-volume of pre-washed streptavidin agarose beads (GE Healthcare Life Science, Cat# 17511301). After 3 h, the beads were transferred to a new 1.5-mL tube and washed once with SDS-Wash buffer (25 mM Tris-HCl, pH 7.4, 2% SDS), twice with RIPA buffer (50 mM Tris-HCl, pH 7.4, 150 mM NaCl, 1 mM EDTA, 1% NP-40, 0.1% SDS, 0.4% sodium deoxycholate), once with TNNE buffer (25 mM Tris-HCl, pH 7.4, 150 mM NaCl, 1 mM EDTA, 0.1% NP-40), and three times with 50 mM ammonium bicarbonate pH 8.0 (ABC buffer). On-bead digestion was performed with 1 µg trypsin (ProMega, Cat# V511A) in 60 µL ABC buffer with shaking overnight at 37°C, followed by further digestion with an additional 0.5 µg trypsin for 3 h. Peptide supernatants were collected in new tubes, the beads were washed twice with MS-grade water, and these supernatants were pooled with the peptide supernatant and subsequently dried by vacuum centrifugation. Peptides were stored at −80°C until MS analysis.

### BioID mass spectrometry

BioID tryptic peptides were resuspended in 24 μL 5% formic acid, and 1/4 of the sample was used each for DDA and DIA on a TripleTOF 6600 MS (SCIEX, Framingham, MA, USA) connected in-line to a 425 Nano-HPLC system (Eksigent Technologies, Dublin, CA, USA). Samples were loaded *via* an autosampler at 800 nL/min (in 0.1% formic acid) onto a home-packed fused silica capillary column (3 μm Reprosil-Pur 120 C18-AQ, Dr. Maisch, Germany; 100 μm internal diameter, 10 cm length) with a 5–8 μm emitter tip opening generated using a laser puller (Sutter Instrument Co, model P-2000, Novato, CA, USA). Peptides were eluted at 400 nL/min in 0.1% formic acid over a 90 min gradient of 2–35% acetonitrile, followed by a 15 min clean-up (80% acetonitrile with 0.1% formic acid) and 15 min equilibration (2% acetonitrile with 0.1% formic acid).

For DDA mode, an MS1 survey scan with 250 ms accumulation time and 400–1800 *m/z* range was followed by up to 10 MS/MS candidate ion scans, each with 100 ms accumulation time and 100–1800 *m/z* range in high sensitivity mode. Only ions with charges of 2+ to 5+ were analyzed, then subsequently excluded for 7 s (with 50 mDa mass tolerance). The isolation width was 0.7 *m/z*, and the minimum threshold was set to 200.

For DIA mode, each 250 ms MS1 scan was followed by 54 variable isolation windows (7–50 Da) covering the 400–1250 amu mass range with a 0.5 Da overlap between acquisition windows. The cycle time was approximately 3.8 s.

DDA and DIA of the same sample were acquired sequentially. Between different samples, two column wash cycles (2 h of alternating 35% acetonitrile with 0.1% formic acid and 80% acetonitrile with 0.1% formic acid), a 30 min bovine serum albumin (BSA) quality control run, and a 30 min BSA mass calibration run (adjusting for mass drift and verifying peak intensity) were performed.

The entire BioID dataset was deposited in ProteomeXchange *via* partner MassIVE (massive.ucsd.edu) and assigned accession numbers PXD075628 and MSV000101129.

### BioID DDA data analysis

MS raw files were stored and searched directly *via* ProHits^56^. Raw files (.WIFF and .WIFF.SCAN) were converted to MGF and mzML formats using ProteoWizard (v3.0.4468)^59^ and AB SCIEX MS Data Converter (V1.3 beta). The database used for searches consisted of the human and adenovirus sequences in the RefSeq protein database (version 57) and mouse protein database (version 53) supplemented with “common contaminants” from the Max Planck Institute (http://141.61.102.106:8080/share.cgi?ssid=0f2gfuB) and the Global Proteome Machine (www.thegpm.org/crap/index.html). The search database consisted of 72,226 entries in total, including forward and reverse sequences (labeled “gi 9999” or “DECOY”).

DDA spectra were analyzed using MSFragger^60^ (*via* FragPipe in ProHits) in label-free quantification mode, using an *in silico* peptide database to mimic trypsin (cleavage at K and R residues with no cuts before P(HF1)) and allowing up to two missed cleavages. No fixed modifications were selected, and methionine oxidation and asparagine/glutamine deamidation were allowed as variable modifications. Precursor and fragment mass tolerances were set to ±20 ppm. Match-between-runs was turned on, with matching permitted to the nearest seven MS runs (to allow matching of peptides across all timepoints within each bait). Normalization between runs and MaxLFQ were turned off. Before each search, automatic mass calibration and parameter optimization were performed independently on each file (including all variable modifications). Other search parameters used the default settings.

Peptide validation was performed *via* the ProteinProphet and PeptideProphet applications of Philosopher^61^ 5.0.0 in ProHits. PeptideProphet was run with the following default closed-search parameters: “--decoyprobs --ppm --accmass --nonparam –expectscore” (and Percolator with min probability 0.5 and settings “--only-psms --no-terminate --post-processing-tdc”), and ProteinProphet was run with “--maxppmdiff 2000000.” This enabled a final protein false discovery rate (FDR) of 1%.

To identify and score high-confidence proximal interactors, the spectral count search results at the protein level were reformatted with an in-house R script for processing by SAINTexpress^62^ 3.6.3-based spectral count analysis *via* the command line. Controls consisted of the empty vector, miniTurbo-3×FLAG, and miniTurbo-3×FLAG-eGFP. We analyzed two independent biological replicates for each condition (five baits and two controls). Default SAINTexpress parameters were used, with no compression of control or bait samples. Each time point was analyzed in a separate SAINTexpress run (with appropriate time point controls), and all results were concatenated into a single file. Preys with BFDRs ≤ 1% were considered high-confidence proximal interactors.

### BioID DIA data analysis

As with the DDA files, DIA raw files were stored and searched *via* ProHits. A spectral library was built from the DDA files using FragPipe (within ProHits) and the same parameters as above, with protein N-terminal acetylation as an additional variable modification). This spectral library was then used to search the DIA files with DIA-NN^63^ 1.8.2 beta 8. Default search parameters were used, except the following: “Robust LC” quantitation strategy, no protein inference, and no MaxLFQ. Proteins were validated with a 1% FDR as described above.

### Candidate selection of differentially localized proteins

To identify preys changing significantly in their proximal association across days, an ANOVA was performed per bait, per prey, on BioID DIA intensities with Benjamini-Hochberg correction for multiple testing correction of p-values. Similarly, ANOVA testing was performed on the total proteome data set to identify significant changes in abundance across days. Coefficients of variation (CVs) were calculated for all proteins in both datasets to check the reproducibility of replicates. The total proteome data set was joined with the BioID DIA intensities, for the 3048 proteins identified across both datasets. Proteins were then filtered for high-confidence proximal interactors, keeping only preys with a SAINT BFDR ≤ 1% in the DDA spectral count analysis (1034 preys with corresponding abundance quantifications). To simplify the distributions and de-emphasize outliers, data were log_2_-transformed after adding 1 to prevent negative values post-transformation. Linear regression (least squares) was performed for each bait, for each prey, to model the relationship between protein abundance and proximal association. Slopes and R^2^ values were extracted from each model. Slopes were adjusted to their absolute value, then log_2_-transformed (after adding 1 to prevent negative values). For each bait, preys were grouped by their adjusted slopes and R^2^ values for their linear models: with 100 cutoff bins for slopes (increasingly stringent from log_2_(slope+1) ≥ 0 up to log_2_(slope+1) ≥ 2)and 50 cutoff bins for R^2^ (increasingly stringent from R^2^ ≥ 0 up to R^2^ = 1), there were with 5,000 combinations of (adjusted) slope-R^2^ pairs in total for each bait.

To evaluate which changes in proximal association were not fully explained by changes in protein abundance, these linear models were compared to a null hypothesis *y* = *x* identity line (where proximity measurements are directly 1:1 proportional to changes in protein abundance). To verify that the null hypothesis was a good estimate of abundance effects on proximity measurements, we fit a linear model, per bait, to all BioID *versus* abundance values (all timepoints together), with a *y*-intercept fixed at 0. The linear model R^2^ ranged from 0.37 to 0.91, with slopes ranging from 0.19 to 0.48. This suggested that a linear model was a moderate-to-excellent fit for the data overall, and a slope of 1 as a null hypothesis was conservative. To compare each bait-prey *versus* abundance relationship to the null hypothesis, an R^2^ value was calculated for each bait-prey pair against a *y* = *x* line to estimate how predictive this identity line would be for the data; we call this the R^2^ identity.

Thresholds for CV, linear model adjusted slope and R^2^, R^2^ identity, and BioID ANOVA adjusted *p*-values were established to facilitate evaluation of prey lists within the slope-R^2^ cutoff bins. The proportions of each list matching these criteria were then averaged to score each bin. Specifically, the five measures calculated were as follows:

1. Proportion of the bin’s list with reproducibility of replicates within 20% CV for each of the BioID and total proteome datasets:

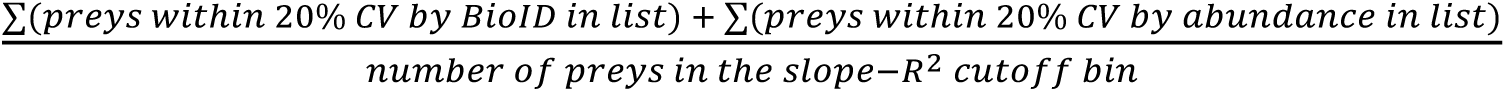

2. Proportion of the list with adjusted slopes within the top quartile for that bait:

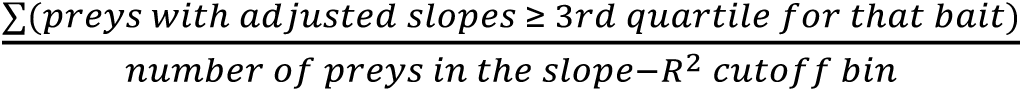

3. Proportion of the list with R2 values within the top quartile for that bait:

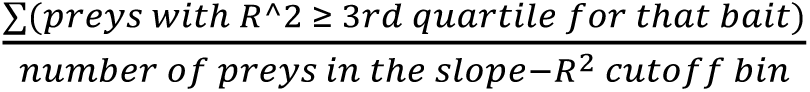

4. Proportion of the list significantly changing in their BioID proximal association across days:

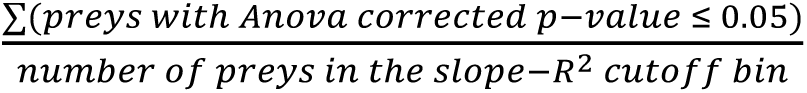

5. Proportion of the list with the lowest R^2^ values against the identity line (R^2^-identity). Note, that R^2^ values calculated against an alternate line than the line of best fit may be negative if using the mean of the values would be a better predictor than the alternate line. In our case, values were indeed predominantly negative (ranging from approximately −10^6^ to 0.4, with even the 3^rd^ quartile around −90). As such, to identify **lowest** R^2^-identity values, we took the absolute value and then log_2_-transformed (adding 1 to prevent negatives), and then leniently selected the 3^rd^ quartile of these values, per bait, to serve as a cutoff for low R^2^-identity. R^2^-identity values higher than these cutoffs were those that were more negative prior to adjustment; hence, we selected preys **above** these R^2^-identity cutoffs as those with the greatest deviation from the identity line:

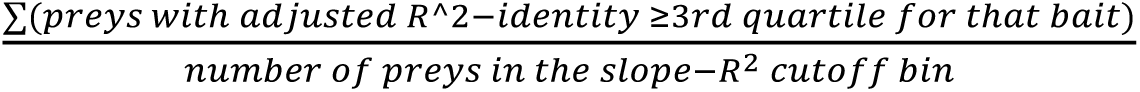

Each of these five measures were averaged across baits, and a mean score across all measures was calculated for each (adjusted) slope-R^2^ cutoff bin. The resulting mean scores were sorted in decreasing order, and the top 2% of the lists were selected. The lowest (adjusted) slope and R^2^ cutoff among these lists were selected, and this combination was used as an ‘extended 2% list’ (slope ≥ 1.47 and R^2^ ≥ 0.69). This list was composed of all preys with SAINT BFDR ≤ 1%, with adjusted slope and R^2^ values for linear models above these thresholds, and a significant difference in proximal association across days. Note, that because there were only two replicates for the BioID dataset and, consequently, low statistical power, the difference in proximal association across days was used as a guide rather than a strict filter. As such, preys with a significant change across days from the BioID ANOVA test were allowed *without* necessitating significance after multiple testing correction. In total, this “extended 2% list” contained 159 unique preys in total across 27 baits.

### siRNA screening

siRNA constructs (SMARTpools and deconvolved pools) were originally purchased from Dharmacon, with SMARTpools (mouse siGenome library) accessed through the NBCC at LTRI. siRNA was transfected (20 nM final) into cells (10,000 cells at 33% reprogramming potential in 48-well format) using Lipofectamine™ RNAiMax (Invitrogen) in Opti-MEM™ (Gibco), per Invitrogen’s recommendations, and reprogramming was induced with Dox for 6 days in triplicate, with fresh mES medium with Dox refreshed daily. Cells were fixed with 4% paraformaldehyde (PFA) and stained for AP (Vector Red; Vector Labs) and DAPI (nuclear counterstaining). Colony formation was imaged on an IN Cell Analyzer 6000 (GE Healthcare) with a 4× objective in both the AP (dsRed) and DAPI channels across four non-overlapping fields per well (initial screen) or on an Incucyte SX5 (Sartorius) with a 4× objective in both the AP (dsRed) and DAPI channels across the whole well (colony formation experiments). For the initial screen, images were analyzed using a custom platform on the Columbus Image Data Storage and Analysis System (PerkinElmer, version 2.3), which measured AP-positive, high-DAPI density areas to define and quantify colonies. The overall colony area in each well was determined using the sum of the overlap area of the four fields captured, log_10_ transformed, and averaged from the three replicates. Complete Columbus scripts are available upon request. For the colony formation experiments, images were analyzed using a custom analysis platform with Incucyte GUI software version 2023A (Sartorius), which also measured AP-positive, high-DAPI density areas to define and quantify colonies. Triplicate colony counts were averaged. Complete Incucyte scripts are available upon request.

### Analysis of siRNA candidates

Statistical significance for differences in AP-positive colony area in the initial screen was calculated by ANOVA followed by Dunnett’s test for multiple comparisons (DescTools package in R, version 0.99.60), against the Dox control. Within each significance bin (*p*-values ≤ 0.0001, 0.001, 0.01, 0.05, and 1), siRNA constructs were ordered by average colony area, log_10_-transformed (adding 1 to prevent negative values), and visualized in a scatterplot using the ggplot2 R package (version 3.3.5). Colony counts were analyzed by one-way ANOVA for significance and visualized using GraphPad Prism version 10.4.0.

### RT-qPCR

Total RNAs were extracted from knockdown and control samples using a RNeasy Mini Kit (Qiagen) per the manufacturer’s instructions. A 500 ng sample of RNA was used for reverse transcription using a High-Capacity cDNA Reverse Transcription kit (Applied Biosystems) with random hexamer primers per the manufacturer’s instructions. Absolute qPCR was performed with the PCR primers in **Supplementary Table S8** using PowerTrack SYBR Green Master Mix for qPCR (Applied Biosystems) on a CFX Opus Real-Time PCR system (Bio-Rad). Each transcript was normalized to the *Actb* level and are graphed relative to the highest value of the sample set shown (set to 100).

### Immunofluorescence

For all immunofluorescence images (transduced, non-transduced, and knockdown sample sets), cell populations were plated on 1% gelatin-coated coverslips. Cells were washed twice with phosphate-buffered saline containing calcium and magnesium (PBS++) and subsequently fixed in 4% PFA in PBS++ for 10 min. Cells were washed with PBS, permeabilized with 0.25% Triton X-100 in PBS for 10 min, and blocked in either 4% BSA or 4% milk powder in PBS for 1 h. Bait proteins were detected using a monoclonal mouse anti-FLAG antibody (M2, Sigma-Aldrich, Cat# F3165; used at 1:1500), CDH1 was detected with a monoclonal rabbit anti-E-cadherin antibody (Cell Signaling Technology, Cat# 24E10; used at 1:2000), OCT4 with a monoclonal mouse anti-OCT3/4 antibody (C-10, Santa Cruz Biotechnology, Cat# sc-5279; used at 1:500), PKP1 with a monoclonal mouse anti-PKP1 antibody (10B2, Santa Cruz Biotechnology, Cat# sc-33636; at 1:500), and JUP with a monoclonal mouse anti-y-catenin antibody (D-12, Santa Cruz Biotechnology, Cat# sc-398183; used at 1:500) in 2% BSA in PBS. Secondary detection was performed using goat anti-mouse Alexa Fluor^TM^ 488 (Molecular Probes, Thermo Fisher Scientific, A11001, used at 1:1000) or goat anti-rabbit Alexa Fluor^TM^ 488 (Molecular Probes, Thermo Fisher Scientific, A11034, used at 1:1000), goat anti-mouse Alexa Fluor^TM^ 594 (Molecular Probes, Thermo Fisher Scientific, A11005, used at 1:1000), or goat anti-rabbit Alexa Fluor^TM^ 594 (Molecular Probes, Thermo Fisher Scientific, A11012, used at 1:1000), goat anti-rabbit Alexa Fluor^TM^ 647 (Molecular Probes, Thermo Fisher Scientific, A21244, used at 1:1000), Alexa Fluor^TM^ 647 phalloidin labeling probe (Molecular Probes, Thermo Fisher Scientific, A22287, used at 1:500), and Alexa Fluor^TM^ 594 streptavidin conjugate (Molecular Probes, Thermo Fisher Scientific, Cat# S11227, used at 1:3000) in 2% BSA in PBS. DAPI (Sigma-Aldrich, 10 mg/mL, used at 1:10,000) was used as a nuclear counterstain. Coverslips were mounted in ProLong Gold AntiFade (Molecular Probes, Thermo Fisher Scientific, Cat# P36930) and imaged on a Nikon C1Si confocal microscope with a 60× oil-immersion lens or on a Nikon Eclipse Ti2 with a 60× oil-immersion lens. Images were processed using EZ-C1 FreeViewer software (v3.90, Nikon) or NIS Elements software (v5.41.02, Nikon).

### Immunoblotting

Cells were lysed as described in *Streptavidin capture and digestion* and an aliquot of clarified lysate was boiled in Laemmli SDS-polyacrylamide gel electrophoresis (PAGE) sample buffer. Proteins were resolved on 10% polyacrylamide gels and transferred to nitrocellulose membranes (GE Healthcare Life Science, Uppsala, Sweden, Cat# 10600001) for immunoblotting. Membranes were either blocked using 5% BSA in Tris-buffered saline with 0.1% Tween-20 (TBST; to identify biotinylated proteins) or 5% milk powder in TBST (to identify bait proteins). Baits were probed using a monoclonal mouse anti-FLAG antibody (M2, Sigma-Aldrich, Cat# F3165; used at 1:2000) in 3% milk in TBST, washed in TBST, and detected with 1:10,000 sheep anti-mouse IgG-horseradish peroxidase (HRP; GE Healthcare Life Science, Cat# NA931) for 1 h. Biotinylated proteins were probed using HRP-conjugated streptavidin (GE Healthcare Life Science, Cat# RPN1231vs) at 1:3000 in BSA blocking buffer for 1 h. A monoclonal mouse anti-glyceraldehyde 3-phosphate dehydrogenase (GAPDH) antibody (G-9, Santa Cruz Biotechnology, Cat# sc-365062) was used at 1:1000 as a loading control. Proteins were visualized using LumiGLO chemiluminescent reagent (Cell Signaling Technology, USA, Cat# 7003S) and imaged using a Bio-Rad ChemiDoc (Bio-Rad, Mississauga, ON, Canada).

### Data visualization and enrichment analysis

Bait-prey dot plots were generated using ProHits-viz^64^ (http://prohits-viz.org). Once a prey passed the ≤1% BFDR threshold with one bait, all its quantitative (intensity) values across all baits shown were recovered with the SAINT BFDR represented by the edge color intensity. The BFDR of the prey with each bait is indicated by the node’s edge color (with a dotted line representing no SAINT information from DDA), and the log_2_-average intensity (from DIA) is represented by the node’s color gradient. The size of the node indicates the relative prey counts across all baits, relative to the bait with which it was detected with the highest number of spectral counts.

GO enrichment analysis was run in Python using the g:Profiler-official package (version 0.3.5) and the mouse database. Enrichment dot plots were generated in Python with a package available on Github (https://github.com/gingraslab/go_terms_dot_plot) that visualizes the top enriched GO terms of gene lists from different clusters (900 max term size, top three terms per condition in a filled manner). Clusters were run both together using a multiquery approach (to assign *p*-values across clusters) and individually (to assign intersections for each term).

Significant prey counts of cell junction terms (by Gene Ontology Cellular Compartment) was determined by the intersection size of the list of high confidence proximity interactors (SAINT BFDR ≤ 0.01) for each indicated GO:CC term for each cell junction-associated bait (by initial bait subcellular localization) and time point. Aggregated counts for each GO:CC term across these baits were merged by day for total interaction counts.

## Acknowledgements

The authors wish to acknowledge the Model Production Core at the Centre for Phenogenomics, Sinai Health, for generating iPSC embryo chimeras, and High-Fidelity Science Communications for manuscript editing. Proteomics and flow cytometry work was performed at the NBCC, a facility supported by Canada Foundation for Innovation funding, the Government of Ontario and Genome Canada and Ontario Genomics (OGI-139). This work was supported by grants from the Canadian Institutes of Health Research Foundation (CIHR FDN143301 and PJT-185987), Terry Fox Research Institute (TFRI; PPG 1107), and the Natural Sciences and Engineering Research Council of Canada (NSERC Discovery Grant RGPIN-2019-06297). Anne-Claude Gingras holds the Canada Research Chair (Tier 1) in Functional Proteomics and the inaugural Lou Siminovitch Sinai 100 Chair. Reuben Samson is supported by an Ontario Graduate Scholarship. Julia Kitaygorodsky is supported by a Canada Graduate Scholarship – Doctoral from the Natural Science and Engineering Research Council of Canada.

## Abbreviations

2° MEF: secondary mouse embryonic fibroblast
AP: alkaline phosphatase
AP-MS: affinity purification–mass spectrometry
DDA: data-dependent acquisition
DIA: data-independent acquisition
Dox: doxycycline
GO: gene ontology
iPSC: induced pluripotent stem cell
LC-MS: liquid chromatography–mass spectrometry
MEF: mouse embryonic fibroblast
MET: mesenchymal-to-epithelial transition
MS: mass spectrometry
mT: miniTurbo
OKMS: OCT4, KLF4, MYC, SOX2 (reprogramming factors)
PDB: proximity-dependent biotinylation
RT-qPCR: reverse transcription-quantitative polymerase chain reaction
rtTA: reverse tetracycline-controlled transactivator
TMT: tandem mass tag

