## Supplementary material for "Spatiotemporal remodeling of cytoskeletal and junction networks during somatic cell reprogramming": supp table legends

| **Table S1** |  |
| --- | --- |
| Reprogramming efficiency of harvested embryo batches | |
| **Sheet descriptions:** |  |
| *Batch Analysis* | Reprogramming potential of harvested and banked batches as measured by flow cytometry |

| **Table S2** |  |
| --- | --- |
| Total proteome dataset |  |
| **Sheet descriptions:** |  |
| *This study* | Total proteome analysis FragPipe output (TMT16 multiplex) |
| *Dataset comparison* | Comparison of identified proteins (as gene names) between this study and two previously published datasets (Benevento 2014 - doi:10.1038/ncomms6613; Hansson 2012 - doi:10.1016/j.celrep.2012.10.014) that used similar 2° MEF reprogramming models |
| *Clustering* | Fuzzy c-means clustering analysis of the total proteome intensities, normalized by Day 0 and standardized between -2 to 2 per protein. Membership scores across clusters are shown for each protein ID, and the top assigned cluster is noted. |
| *GO* | Gene ontology (GO) enrichment analysis with g:Profiler for each cluster of the total proteome dataset (multiquery). Cluster composition is based on the top assigned cluster per protein ID. |

| **Table S3** |  |
| --- | --- |
| Bait selection and expression levels |  |
| **Sheet descriptions:** |  |
| *Bait Description* | Description of selected bait proteins, localizing to cytoskeletal and junctional compartments, chosen based on their stable expression in the total proteome dataset |
| *Total proteome expression* | Intensity fold changes for selected baits across all studied time points; paired t-test results, and coefficients of variation (CVs) are noted |
| *BioID bait expression* | Peptide-level quantification of the miniTurbo enzyme from the BioID dataset as an estimate of bait expression |

| **Table S4** |  |
| --- | --- |
| Proximity-dependent biotinylation (BioID) dataset | |
| **Sheet descriptions:** |  |
| *BioID DDA* | DDA spectral count and intensity quantification for all prey proteins recovered in the BioID dataset (full FragPipe output) |
| *BioID DIA* | DIA intensity quantification for all prey proteins recovered in the BioID dataset (full FragPipe output) |
| *DDA spc SAINT* | Significance scoring by SAINTexpress (spectral count-based analysis) for identified proteins; concatenated SAINT files from each time point analysis. Three additional columns are appended on the right for the: DDA intensity values, average intensity across replicates, and the intensities across the controls. |
| *SAINT combined with DIA* | SAINT file (DDA spc) combined with the DIA intensities. A placeholder BFDR of 1 was used for any bait-prey pairs seen only in the DIA dataset (and thus not scored by SAINT for the DDA analysis) |
| *BioGrid benchmarking* | Benchmarking of each bait's high-confidence (BFDR ≤ 1%) preys against human orthologs in BioGRID |
| *Clustering* | Hierarchical clustering of preys by their BioID DIA intensities, filtered for preys below a 1% BFDR threshold by SAINT (for at least one time point). The resulting prey dendrogram was divided into 15 clusters. |
| *GO* | Multiquery GO analysis for the 15 prey clusters. |

| **Table S5** |  |
| --- | --- |
| Differential associations of cytoskeletal and cell-cell junction regulators during reprogramming | |
| **Sheet descriptions:** |  |
| *CTNNA1 preys* | Multiquery GO analysis for significant preys identified by the CTNNA1 bait for each time point |
| *Cell junctions* | Number of significant preys localizing to the cellular junction components for selected baits and days (corresponds to Figure 3F) |
| *Cornified envelope abundances* | Scaled abundances (from total proteome quantification; scaled from 0–1) for proteins associated with the cornified envelope, and identified by BioID with KRT7 as bait |
| *Desmosome abundances* | Scaled abundances (from total proteome quantification; scaled from 0–1) for proteins associated with the desmosome, and identified by BioID with KRT7 as bait |

| **Table S6** |  |
| --- | --- |
| Abundance normalization for candidate selection | |
| **Sheet descriptions:** |  |
| *Abundance-BioID* | Overlap between the total proteome and BioID datasets, joined by UniProt ID, across four early time points of reprogramming (Days 1, 2, 4, 6) |
| *Global Linear Reg* | Global linear regression analysis across all preys (per bait), with a fixed y-intercept of 0; slopes and R2 values for overall linear model fit |
| *Individual Linear Reg* | Linear models fit per bait-prey pair comparing protein abundance to BioID intensity; slopes, R2 values, linear equations, significance of BioID change across time by ANOVA (overall adjusted p-value by Benjamini-Hochberg), BioID CV across time points, total proteome abundance CV across time points, and R2-identity |
| *Evaluation* | Evaluation table, sorted by decreasing mean score, of slope-R2 bins |
| *Selected Bait-Prey Pairs* | From the top 2% of bins (by mean score), the lowest slope and R2 thresholds were selected; these bins were collapsed to yield 205 bait-prey pairs |
| *Final candidates* | Unique list of 159 protein candidates, collapsed across all baits |

| **Table S7** |  |
| --- | --- |
| Stringency evaluation of computational candidate selection method | |
| **Sheet descriptions:** |  |
| *Top100 BioID* | Top 100 unique preys, ranked by the magnitude of their BioID slope across time; bait of origin, linear model details, selected or unselected |

| **Table S8** |  |
| --- | --- |
| siRNA analysis of selected candidates | |
| **Sheet descriptions:** |  |
| *Screen candidates* | Filtering of selected candidates, removing essential genes and focusing on candidates with available siRNA |
| *Selected candidate descriptions* | Description of selected siRNA |
| *siRNA screen* | Results of siRNA screen: ANOVA (and Dunnett test) of AP-positive colony area, morphology |
| *GO* | Gene ontology (GO) enrichment analysis with g:Profiler for knockdowns showing abnormal morphologies or low colony areas |
| *Primers* | Primer information for qPCR experiments |
| *siRNA validation* | RT-qPCR results of selected 8 siRNA follow-up candidates |

| **Table S9** |  |
| --- | --- |
| Validation of reprogramming disruption effects | |
| **Sheet descriptions:** |  |
| *D6,D9 Dox removal* | Colony counts at Day 15 after siRNA and Dox removal at Day 6 and 9 |
| *D12 Dox removal* | Colony counts at Days 9, 12 and 21 after siRNA and Dox removal at Day 12 |
| *RT-qPCR* | RT-qPCR results for MET and reprogramming/pluripotency markers |
