## Supplementary material for "Spatiotemporal remodeling of cytoskeletal and junction networks during somatic cell reprogramming": Supp Figures

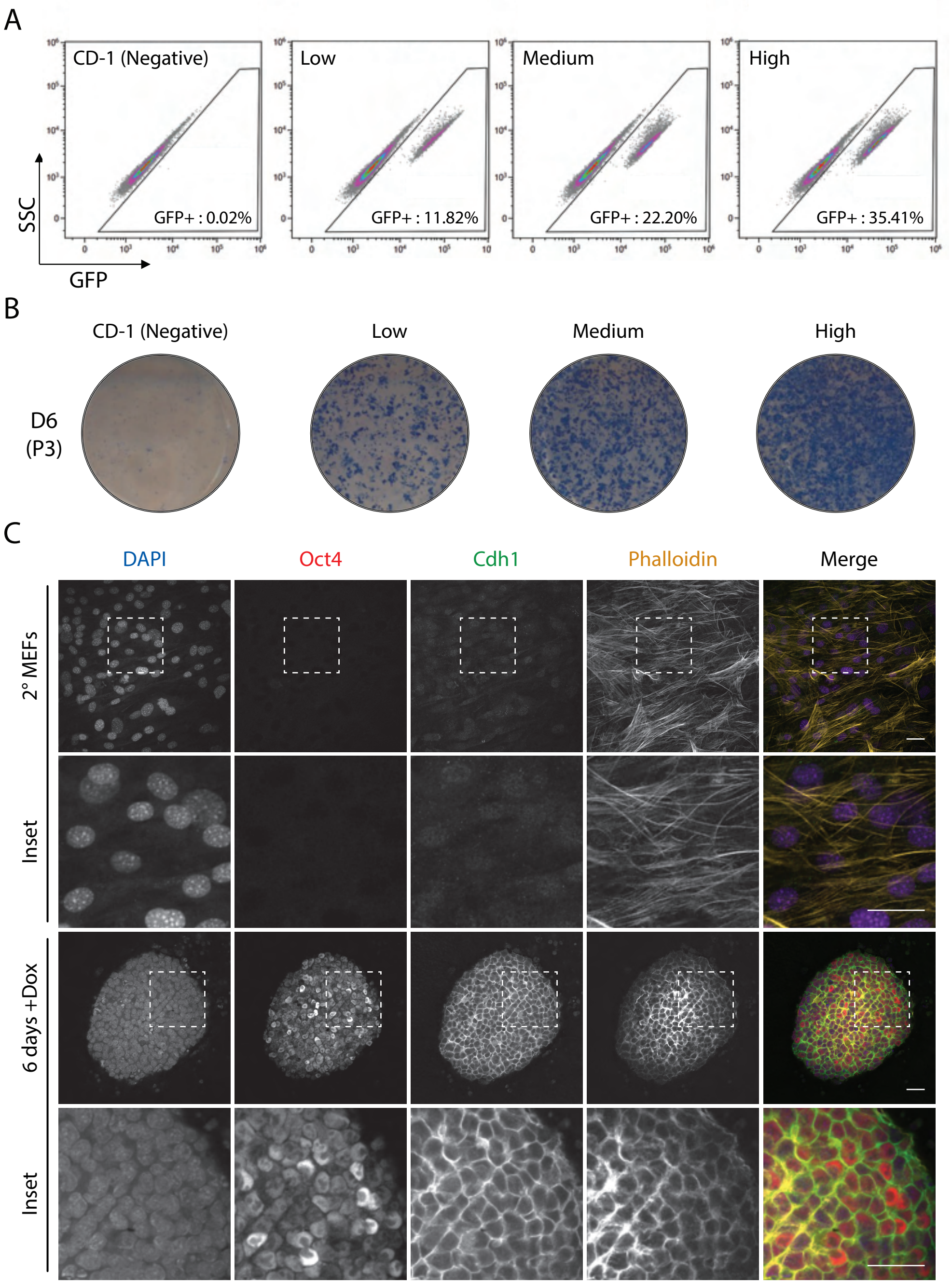
A
CD-1 (Negative)
GFP+ : 0.02%
Low
GFP+ : 11.82%
Medium
GFP+ : 22.20%
High
GFP+ : 35.41%
SSC
GFP
B
CD-1 (Negative)
Low
Medium
High
D6
(P3)
C
DAPI
Oct4
Cdh1
Phalloidin
Merge
2° MEFs
Inset
6 days +Dox
Inset

**Figure S1. Generation of secondary reprogramming mouse fibroblast populations** (**A**) Four large batches of secondary reprogramming mouse fibroblasts with varying reprogramming efficiencies were isolated and pooled from individual chimeric mouse embryos as previously described[28]. Batches were verified for reprogramming potential (GFP positivity) by fluorescence-activated cell sorting analysis and (**B**) assessed for initiation of reprogramming by alkaline phosphatase (AP) staining 6 days post-doxycycline treatment. (**C**) Untreated secondary mouse embryonic fibroblasts (2° MEFs) and those treated with Dox for 6 days (6 days+Dox) were fixed and stained for Oct4 (red), Cdh1 (green), F-actin (Phalloidin, yellow), and nuclei (DAPI, blue) and imaged by confocal microscopy. Scale bars: 20 µm.

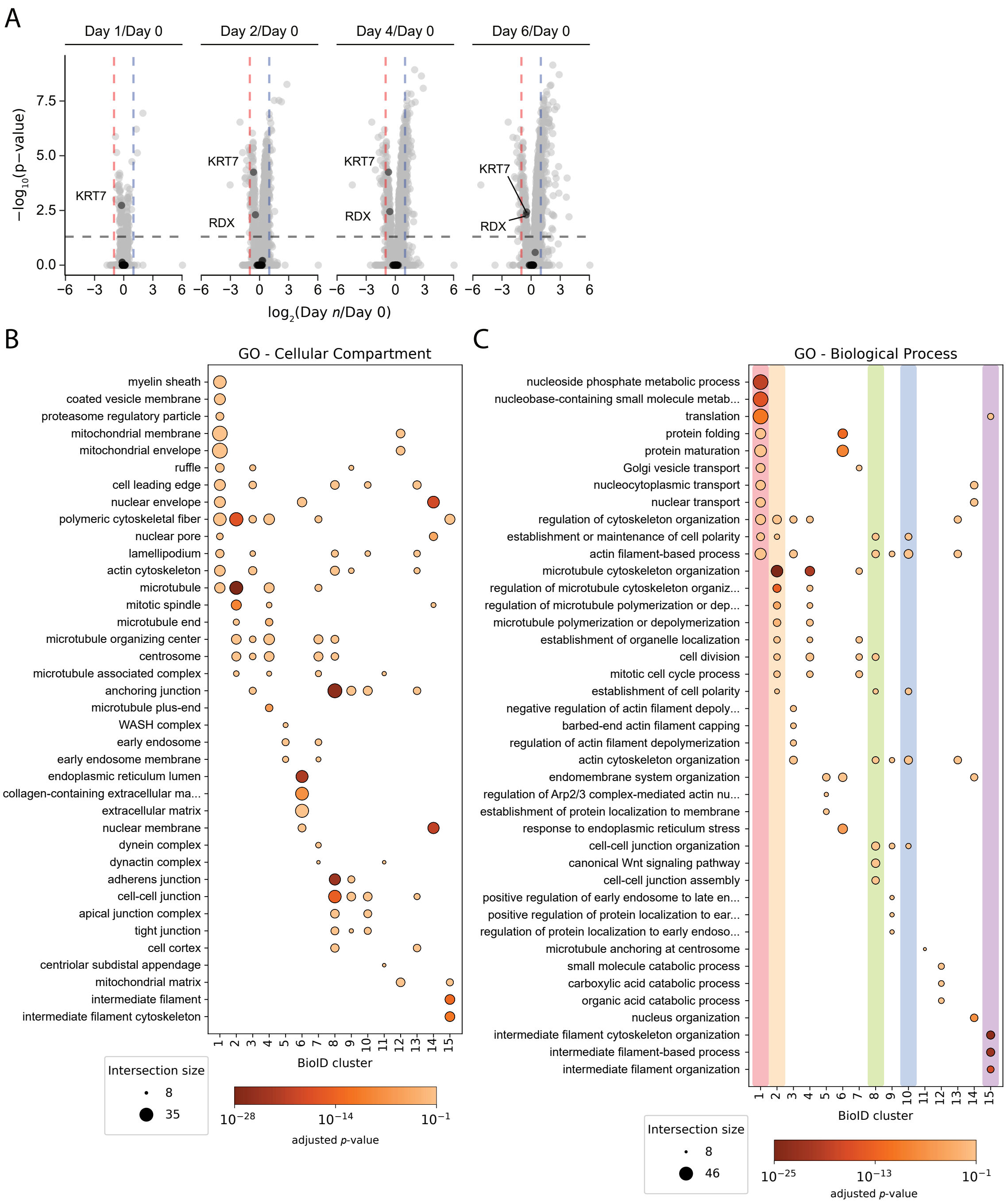
A
Day 1/Day 0
Day 2/Day 0
Day 4/Day 0
Day 6/Day 0
KRT7
RDX
−log10(p−value)
log2(Day n/Day 0)
B
GO - Cellular Compartment
myelin sheath
coated vesicle membrane
proteasome regulatory particle
mitochondrial membrane
mitochondrial envelope
ruffle
cell leading edge
nuclear envelope
polymeric cytoskeletal fiber
nuclear pore
lamellipodium
actin cytoskeleton
microtubule
mitotic spindle
microtubule end
microtubule organizing center
centrosome
microtubule associated complex
anchoring junction
microtubule plus-end
WASH complex
early endosome
early endosome membrane
endoplasmic reticulum lumen
collagen-containing extracellular ma...
extracellular matrix
nuclear membrane
dynein complex
dynactin complex
adherens junction
cell-cell junction
apical junction complex
tight junction
cell cortex
centriolar subdistal appendage
mitochondrial matrix
intermediate filament
intermediate filament cytoskeleton
BioID cluster
Intersection size
8
35
adjusted p-value
C
GO - Biological Process
nucleoside phosphate metabolic process
nucleobase-containing small molecule metab...
translation
protein folding
protein maturation
Golgi vesicle transport
nucleocytoplasmic transport
nuclear transport
regulation of cytoskeleton organization
establishment or maintenance of cell polarity
actin filament-based process
microtubule cytoskeleton organization
regulation of microtubule cytoskeleton organiz...
regulation of microtubule polymerization or dep...
microtubule polymerization or depolymerization
establishment of organelle localization
cell division
mitotic cell cycle process
establishment of cell polarity
negative regulation of actin filament depoly...
barbed-end actin filament capping
regulation of actin filament depolymerization
actin cytoskeleton organization
endomembrane system organization
regulation of Arp2/3 complex-mediated actin nu...
establishment of protein localization to membrane
response to endoplasmic reticulum stress
cell-cell junction organization
canonical Wnt signaling pathway
cell-cell junction assembly
positive regulation of early endosome to late en...
positive regulation of protein localization to ear...
regulation of protein localization to early endoso...
microtubule anchoring at centrosome
small molecule catabolic process
carboxylic acid catabolic process
organic acid catabolic process
nucleus organization
intermediate filament cytoskeleton organization
intermediate filament-based process
intermediate filament organization
BioID cluster
Intersection size
8
46
adjusted p-value

**Figure S2. Selection and enrichment analyses of bait profiles** (**A**) Protein intensities were compared using paired *t*-tests, with *p*-values adjusted using the Bonferroni correction for multiple testing. Each panel shows the $\log_2$(fold-change) of protein abundance between the indicated days (x-axis) and the $-\log_{10}$(adjusted *p*-value) (y-axis). Black points represent chosen bait proteins, which were largely unchanged across conditions. Fold-change (±2-fold) and significance (adjusted $p$-value $\leq 0.05$) thresholds are indicated by dashed lines. (**B**) and (**C**) Dot plots of GO enrichment analysis of proteins in the 15 BioID clusters identified (in Figure 2D) during early reprogramming, restricted to the top three GO terms per cluster (based on the lowest *p*-value; term size limited to 900).

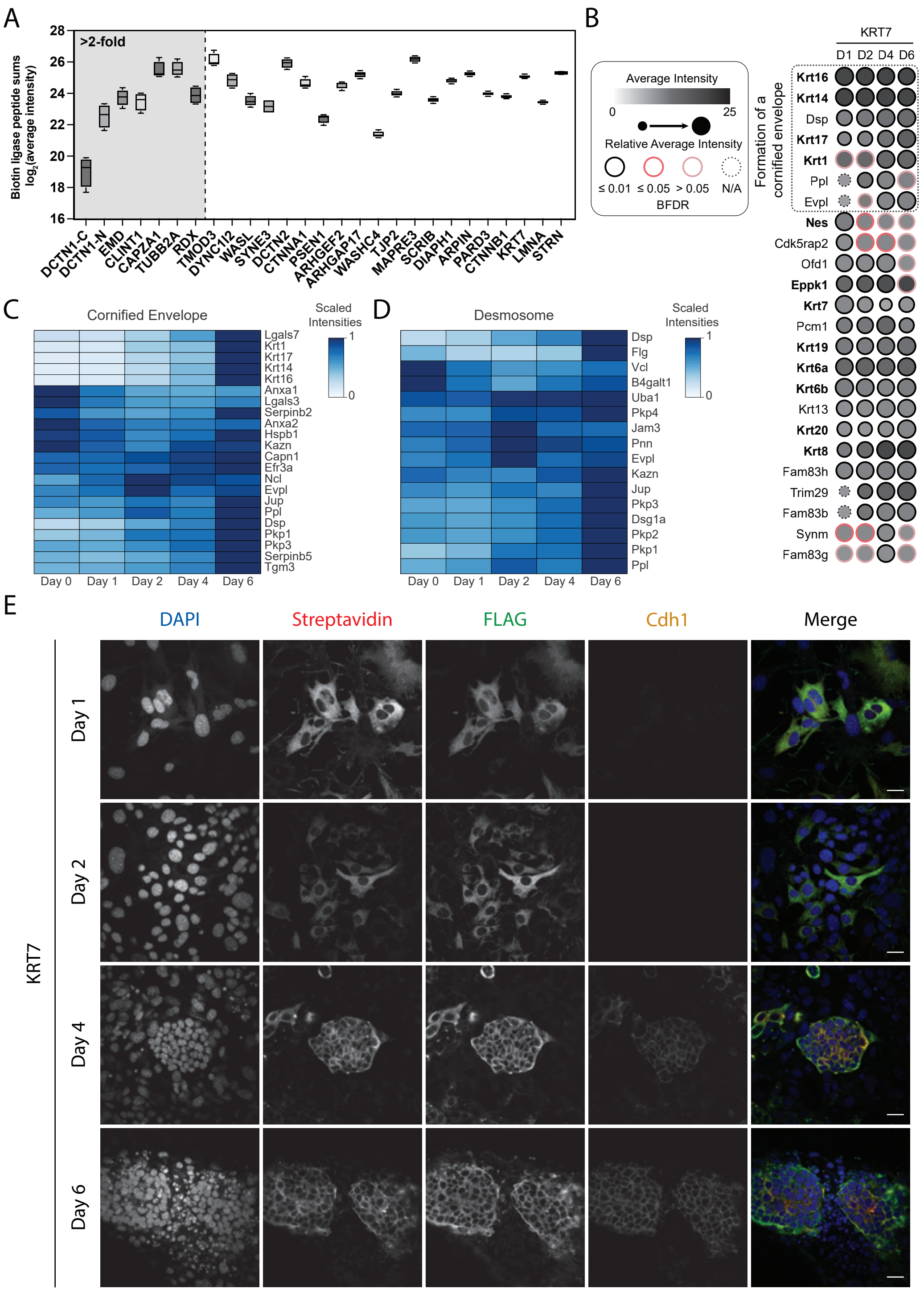
A
B
C
D
E
>2-fold
Biotin ligase peptide sums log2(average intensity)
28
26
24
22
20
18
16
DCTN1-C
DCTN1-N
EMD
CLINT1
CAPZA1
TUBB2A
RDX
TMOD3
DYNC1I2
WASL
SYNE3
DCTN2
CTNNA1
PSEN1
ARHGEF2
ARHGAP17
WASHC4
TJP2
MAPRE3
SCRIB
DIAPH1
ARPIN
PARD3
CTNNB1
KRT7
LMNA
STRN
Average Intensity
0
25
Relative Average Intensity
≤ 0.01
≤ 0.05
> 0.05
N/A
BFDR
KRT7
D1
D2
D4
D6
Formation of a cornified envelope
Krt16
Krt14
Dsp
Krt17
Krt1
Ppl
Evpl
Nes
Cdk5rap2
Ofd1
Eppk1
Krt7
Pcm1
Krt19
Krt6a
Krt6b
Krt13
Krt20
Krt8
Fam83h
Trim29
Fam83b
Synm
Fam83g
Cornified Envelope
Scaled Intensities
1
0
Lgals7
Krt1
Krt17
Krt14
Krt16
Anxa1
Lgals3
Serpinb2
Anxa2
Hspb1
Kazn
Capn1
Efr3a
Ncl
Evpl
Jup
Ppl
Dsp
Pkp1
Pkp3
Serpinb5
Tgm3
Day 0
Day 1
Day 2
Day 4
Day 6
Desmosome
Scaled Intensities
1
0
Dsp
Flg
Vcl
B4galt1
Uba1
Pkp4
Jam3
Pnn
Evpl
Kazn
Jup
Pkp3
Dsg1a
Pkp2
Pkp1
Ppl
Day 0
Day 1
Day 2
Day 4
Day 6
DAPI
Streptavidin
FLAG
Cdh1
Merge
KRT7
Day 1
Day 2
Day 4
Day 6

**Figure S3. Generation of a cornified envelope is supported by BioID.** (**A**) Box plot of total biotin ligase peptides (average intensity, $log_2$-transformed) for each bait across days. Baits with > 2-fold expression level changes across any two time points are indicated with grey shading. (**B**) Dot plot of KRT7-miniTurbo preys across days 1–6. (**C**) Total proteome abundance of proteins that localize to the cornified envelope and (**D**) the desmosome across early reprogramming. Protein intensities are scaled from 0 to 1. (**E**) KRT7-miniTurbo-expressing cells undergoing reprogramming were fixed and stained on days 1, 2, 4, and 6. DAPI (blue), nucleus; streptavidin (red), biotinylation; FLAG (green), miniTurbo-FLAG-tagged bait; Cdh1 (yellow), cell junction. Scale bars: 20 μm.

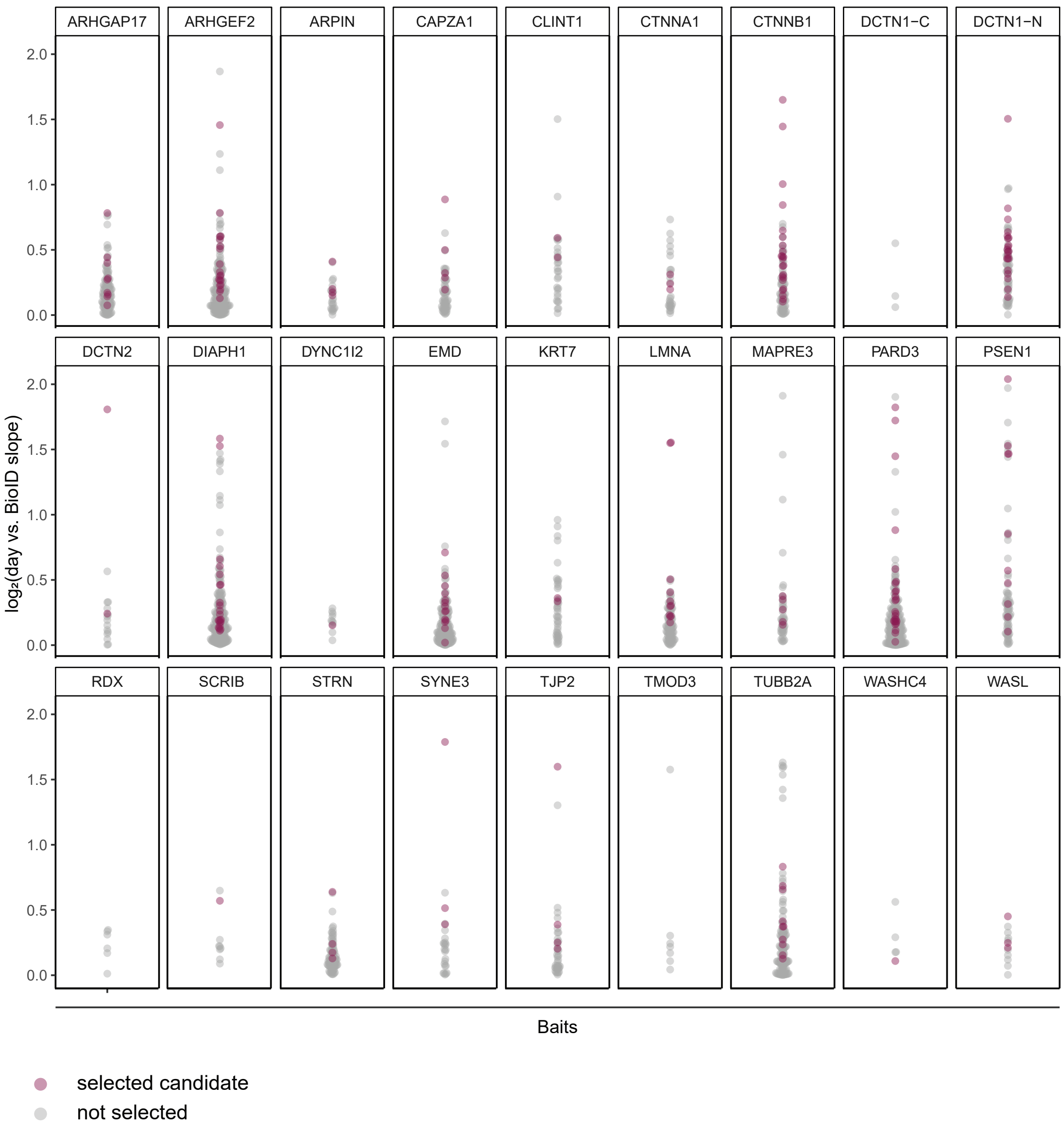
ARHGAP17
ARHGEF2
ARPIN
CAPZA1
CLINT1
CTNNA1
CTNNB1
DCTN1−C
DCTN1−N
DCTN2
DIAPH1
DYNC1I2
EMD
KRT7
LMNA
MAPRE3
PARD3
PSEN1
RDX
SCRIB
STRN
SYNE3
TJP2
TMOD3
TUBB2A
WASHC4
WASL
log₂(day vs. BioID slope)
2.0
1.5
1.0
0.5
0.0
Baits
selected candidate
not selected

**Figure S4. Distribution of siRNA screen candidates among the prey profiles of each bait.** Selected protein candidates for siRNA screening across all baits using the criteria shown in Figure 4G. Only preys with SAINT BFDR ≤ 0.01 are shown. -C and -N indicate C- and N-terminal miniTurbo tags, respectively.

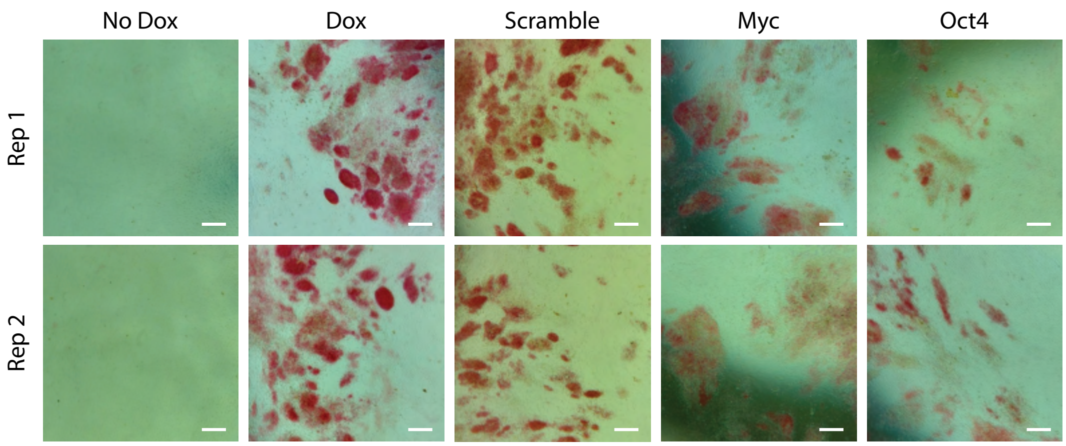

No Dox
Dox
Scramble
Myc
Oct4
Rep 1
Rep 2

**Figure S5. siRNA screen control morphologies** (**A**) Representative images of siRNA screen controls. Red cells indicate alkaline phosphatase positivity. Scale bars: 400 µm.

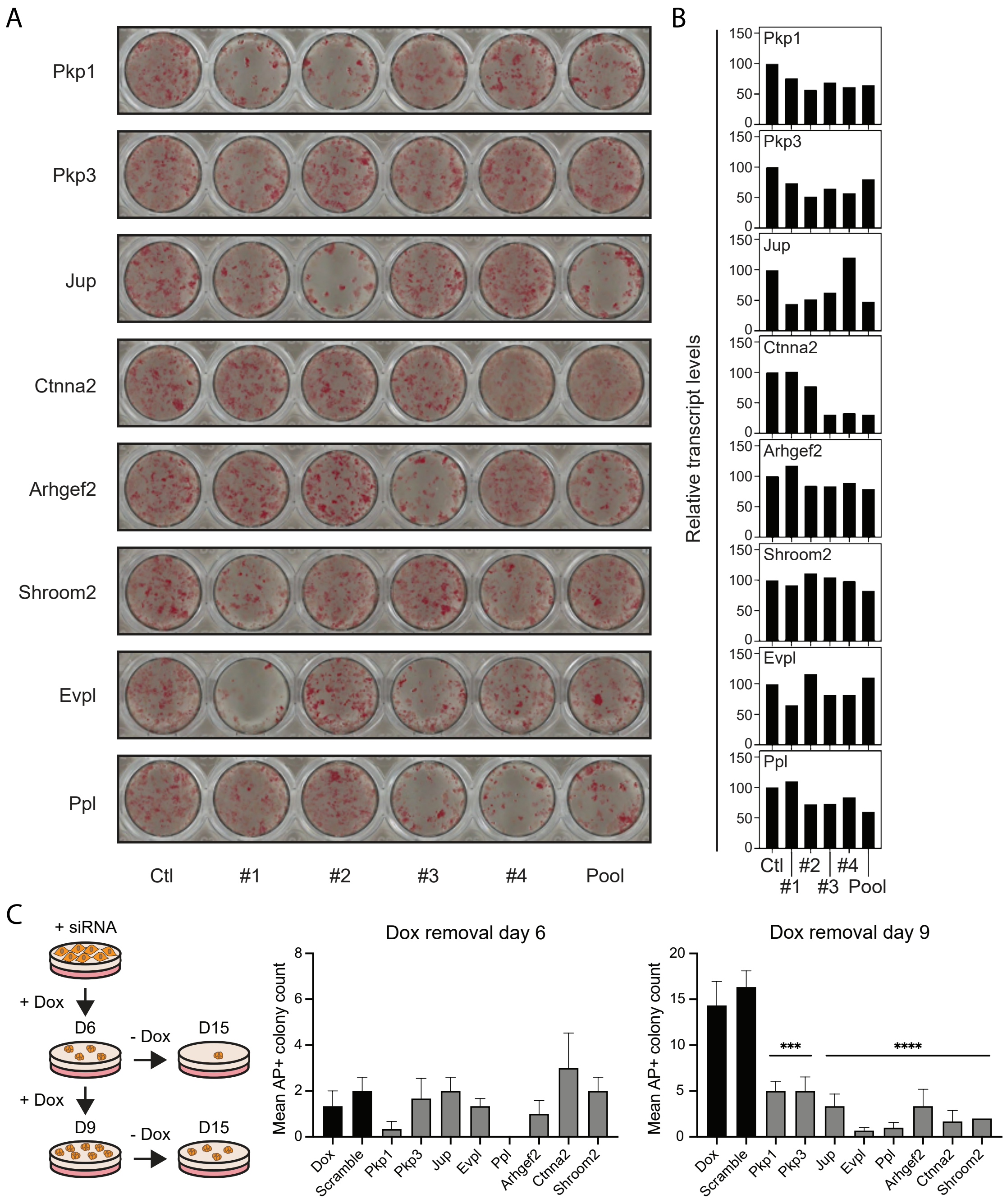
A
Pkp1
Pkp3
Jup
Ctnna2
Arhgef2
Shroom2
Evpl
Ppl
Ctl
#1
#2
#3
#4
Pool
B
Relative transcript levels
Pkp1
Pkp3
Jup
Ctnna2
Arhgef2
Shroom2
Evpl
Ppl
Ctl
#1
#2
#3
#4
Pool
C
+ siRNA
+ Dox
D6
- Dox
D15
+ Dox
D9
- Dox
D15
Dox removal day 6
Mean AP+ colony count
Dox
Scramble
Pkp1
Pkp3
Jup
Evpl
Ppl
Arhgef2
Ctnna2
Shroom2
Dox removal day 9
Mean AP+ colony count
***
****
Dox
Scramble
Pkp1
Pkp3
Jup
Evpl
Ppl
Arhgef2
Ctnna2
Shroom2

**Figure S6. Validation of selected siRNA screen candidates** (**A**) Cells were transfected with individual (#1, #2, #3, #4) or pooled (Pool) siRNAs targeting candidate genes or a scrambled control siRNA (Ctl) and treated with doxycycline (Dox) for 6 days. Alkaline phosphatase (AP) staining was used to detect reprogramming colonies. (**B**) RT-qPCR analysis of siRNA-mediated knockdown efficiency in the samples shown in A. Transcript expression levels were normalized to beta-actin mRNA and measured relative to control samples. (**C**) AP-positive colonies were counted on day 15 after Dox treatment for either 6 or 9 days followed by Dox removal. Asterisks indicate one-way ANOVA adjusted *p*-values (*** = <0.001, **** = <0.0001).

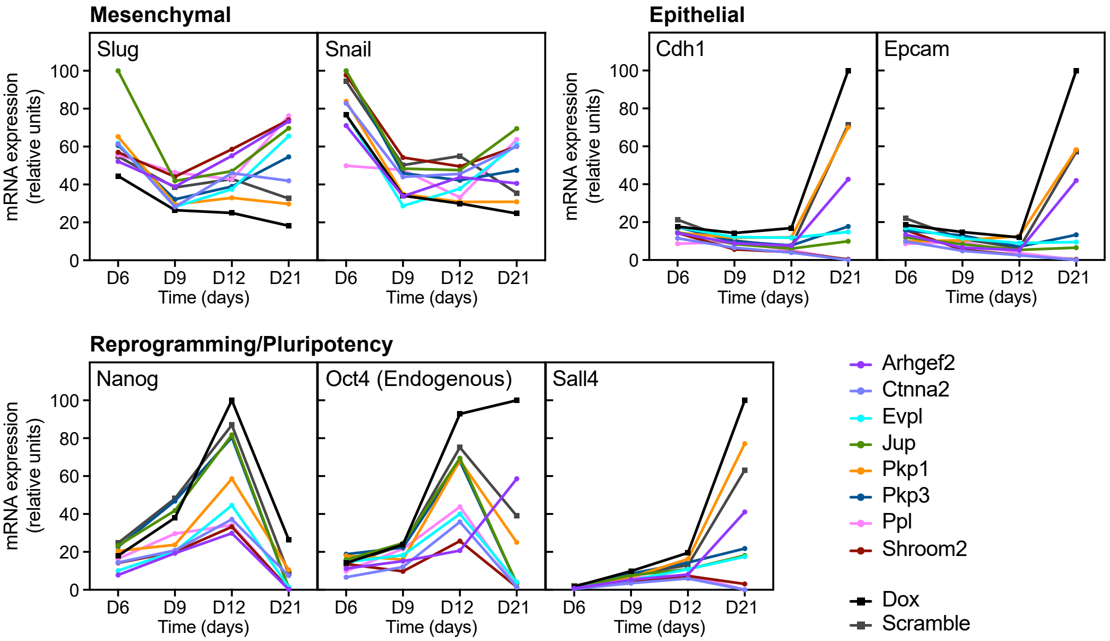
Mesenchymal
Slug
Snail
Epithelial
Cdh1
Epcam
Reprogramming/Pluripotency
Nanog
Oct4 (Endogenous)
Sall4
mRNA expression
(relative units)
Time (days)
D6
D9
D12
D21
0
20
40
60
80
100
Arhgef2
Ctnna2
Evpl
Jup
Pkp1
Pkp3
Ppl
Shroom2
Dox
Scramble

**Figure S7. RT-qPCR of candidate knockdowns** RT-qPCR was performed on mesenchymal, epithelial, and reprogramming/pluripotency markers on days 6, 9, 12, and 21 after siRNA knockdown of the candidates shown in Figure 6A (the Jup, Pkp1, and control data are also shown in Figure 6B). Transcript levels are normalized to beta-actin.
